# Feather aerodynamics suggest importance of lift and flow predictability over drag minimization

**DOI:** 10.1101/2024.05.27.596009

**Authors:** Frida Alenius, Johan Revstedt, L. Christoffer Johansson

**Affiliations:** Department of Biology, Lund University, Lund, Sweden; Department of Energy Science, Lund University, Lund, Sweden

## Abstract

Partly overlapping feathers form a large part of birds’ wing surfaces, but in many species the outermost feathers split, making each feather function as an independent wing. These feathers are complex structures that evolved to fulfil both aerodynamic and structural functions. Yet relatively little is known about how the profile shape and microstructures of feathers impact aerodynamic performance. Here we determined, using fluid dynamic modelling, the aerodynamic capabilities of a section of the primary flight feather forming the leading edge of the split wing tip of a Jackdaw (*Corvus monedula*). Our findings demonstrate that the feather section exhibits a relatively high performance, with lift comparable to manmade aerofoils, however, there is a drag penalty associated with the feather shaft. The model’s vortex shedding behaviour shows low amplitude temporal fluctuations in lift, compared to manmade aerofoils. Notably, the aerodynamic pitch torque around the shaft varies with angle of attack. This, when combined with the built-in pitch-up twist of the feather implies a passive pitch control mechanism for the feather. Taken together, our findings suggest evolutionary adaptations of the flow around the feather, which could be of interest when designing micro-air vehicles and wind turbines.

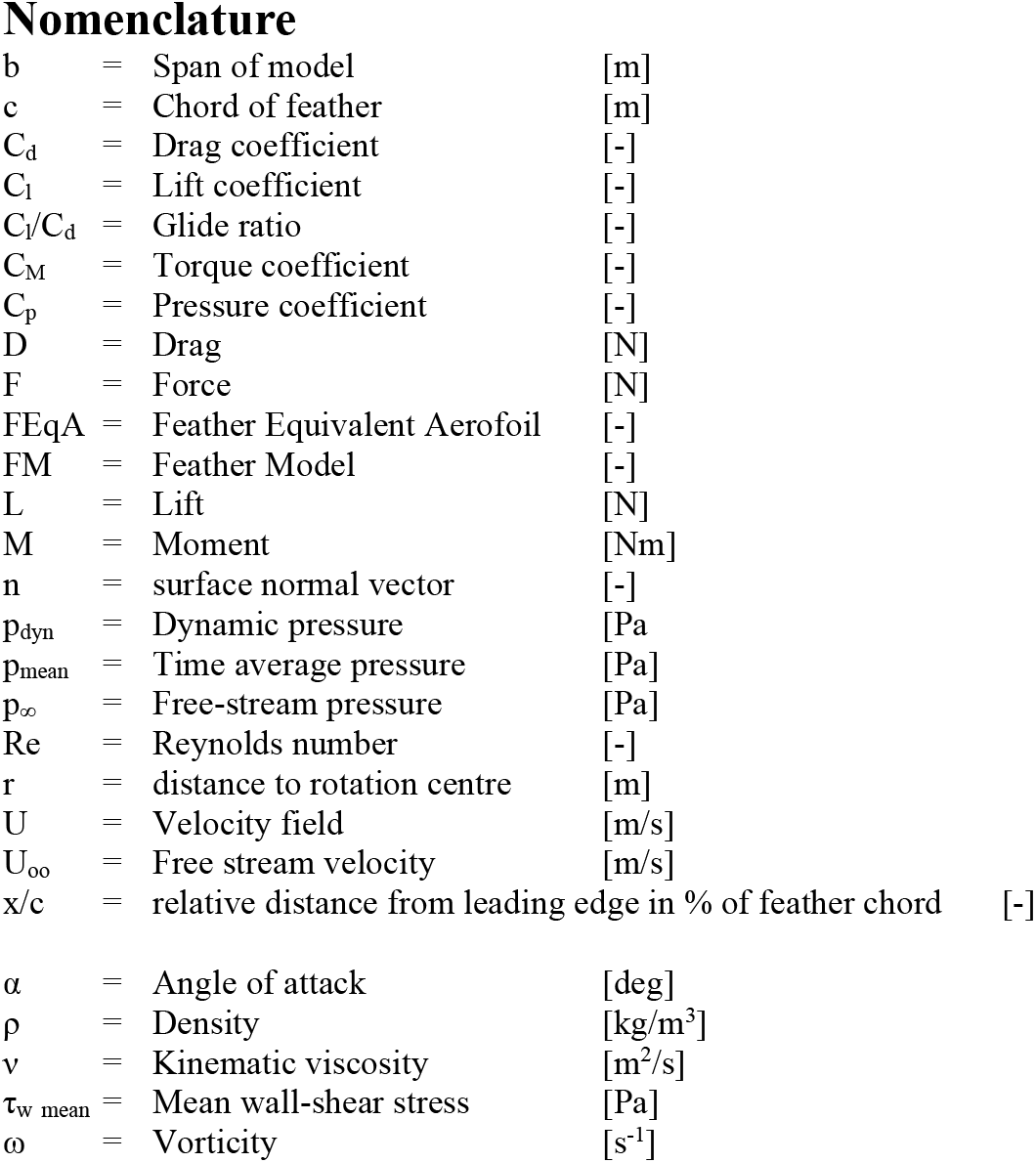

23 Nomenclature
|  |  |  |  |  |
| --- | --- | --- | --- | --- |
| 24 | $b$ | = | Span of model | [m] |
| 25 | $c$ | = | Chord of feather | [m] |
| 26 | $C_d$ | = | Drag coefficient | [-] |
| 27 | $C_l$ | = | Lift coefficient | [-] |
| 28 | $C_l/C_d$ | = | Glide ratio | [-] |
| 29 | $C_M$ | = | Torque coefficient | [-] |
| 30 | $C_p$ | = | Pressure coefficient | [-] |
| 31 | $D$ | = | Drag | [N] |
| 32 | $F$ | = | Force | [N] |
| 33 | FEqA | = | Feather Equivalent Aerofoil | [-] |
| 34 | FM | = | Feather Model | [-] |
| 35 | $L$ | = | Lift | [N] |
| 36 | $M$ | = | Moment | [Nm] |
| 37 | $n$ | = | surface normal vector | [-] |
| 38 | $p_{dyn}$ | = | Dynamic pressure | [Pa] |
| 39 | $p_{mean}$ | = | Time average pressure | [Pa] |
| 40 | $p_{\infty}$ | = | Free-stream pressure | [Pa] |
| 41 | $Re$ | = | Reynolds number | [-] |
| 42 | $r$ | = | distance to rotation centre | [m] |
| 43 | $U$ | = | Velocity field | [m/s] |
| 44 | $U_{\infty}$ | = | Free stream velocity | [m/s] |
| 45 | $x/c$ | = | relative distance from leading edge in % of feather chord | [-] |
| 46 |  |  |  |  |
| 47 | $\alpha$ | = | Angle of attack | [deg] |
| 48 | $\rho$ | = | Density | [kg/m<sup>3</sup>] |
| 49 | $\nu$ | = | Kinematic viscosity | [m<sup>2</sup>/s] |
| 50 | $\tau_{w \text{ mean}}$ | = | Mean wall-shear stress | [Pa] |
| 51 | $\omega$ | = | Vorticity | [s<sup>-1</sup>] |
| 52 |  |  |  |  |
| 53 |  |  |  |  |

## 1 Introduction

Among animals capable of flight, the wings of birds stand out by being built from multiple, partly overlapping, individual structures – feathers. The fact that feathers are individual structures allows the outermost primary flight feathers in many birds to separate and function as aerofoils on their own ^1^, as seen in the multi-slotted wingtips (Figure 1) of, for example, birds of prey ^2^. The morphology of these latter feathers shows distinct properties, such as a smaller chord in the part that work as an independent aerofoil than in the part where feathers overlap as well as a high asymmetry in the location of the shaft along the chord ^3^. Although these feathers have drawn scientific interest ^3–5^, relatively few studies have tested how they perform aerodynamically, and then not in great detail ^1,4,6^.

**Figure 1.**
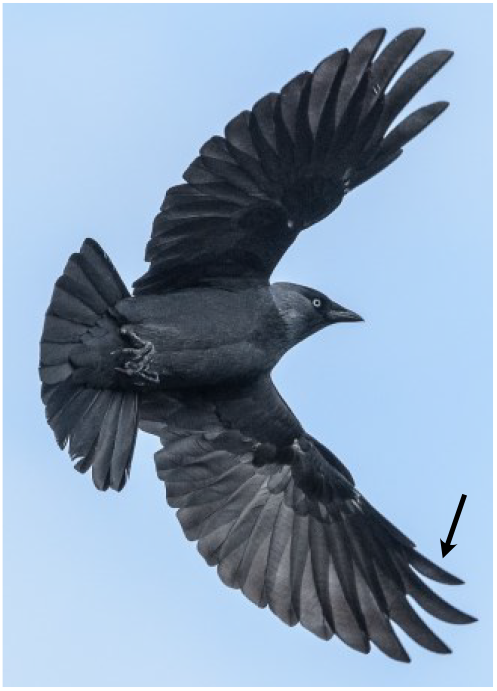
Jackdaws (*Corvus monedula*) have wings where the slotted outer primary feathers can spread out. The arrow points to the ninth primary feather, which is the leading feather in the slotted outer wing and used to create our model. Photo: Arend Vermazeren, used under a Creative Commons licence 2.0

Feathers are very complex structures, where each feather consists of the same structural elements; shaft, barbs and barbules. The barbs branch off at an angle from the shaft and the barbules branch off from the barbs ^3,5,7^ (Figure 2). The composition, size, strength and interaction of the different feather structural elements differ between feather types ^3,7^. A primary flight feather, the feather type of interest here, has two asymmetrical vanes with the leading vane (lateral) being narrower than the trailing vane (medial) ^3,7^. The vanes are built from the barbs and barbules where the barbules are arranged in a hook and groove system which connects to the barbules of adjacent barbs (Figure 2), which creates a cohesive aerodynamic surface ^3,5,7,8^. During flight, the feather bend, twist and sweep in a complex manner ^9^, where the shaft transfers the forces from the barbs to the wing and the body. Hence, the feather structures must be strong enough to transfer the forces, to the body, without breaking or buckling, e.g. by having a sufficient diameter. Since multiple functions have to coexist in the feather, adaptations for flight could have led to compromises and less than optimal features compared to if only one function was considered ^10^. The goal of generating forces efficiently and maintaining structural integrity may thus not always align, and we do not, for example, know how the shaft, extending below the lower surface of the barbs (likely for structural reasons), effect the aerodynamic properties of the feather.

**Figure 2.**
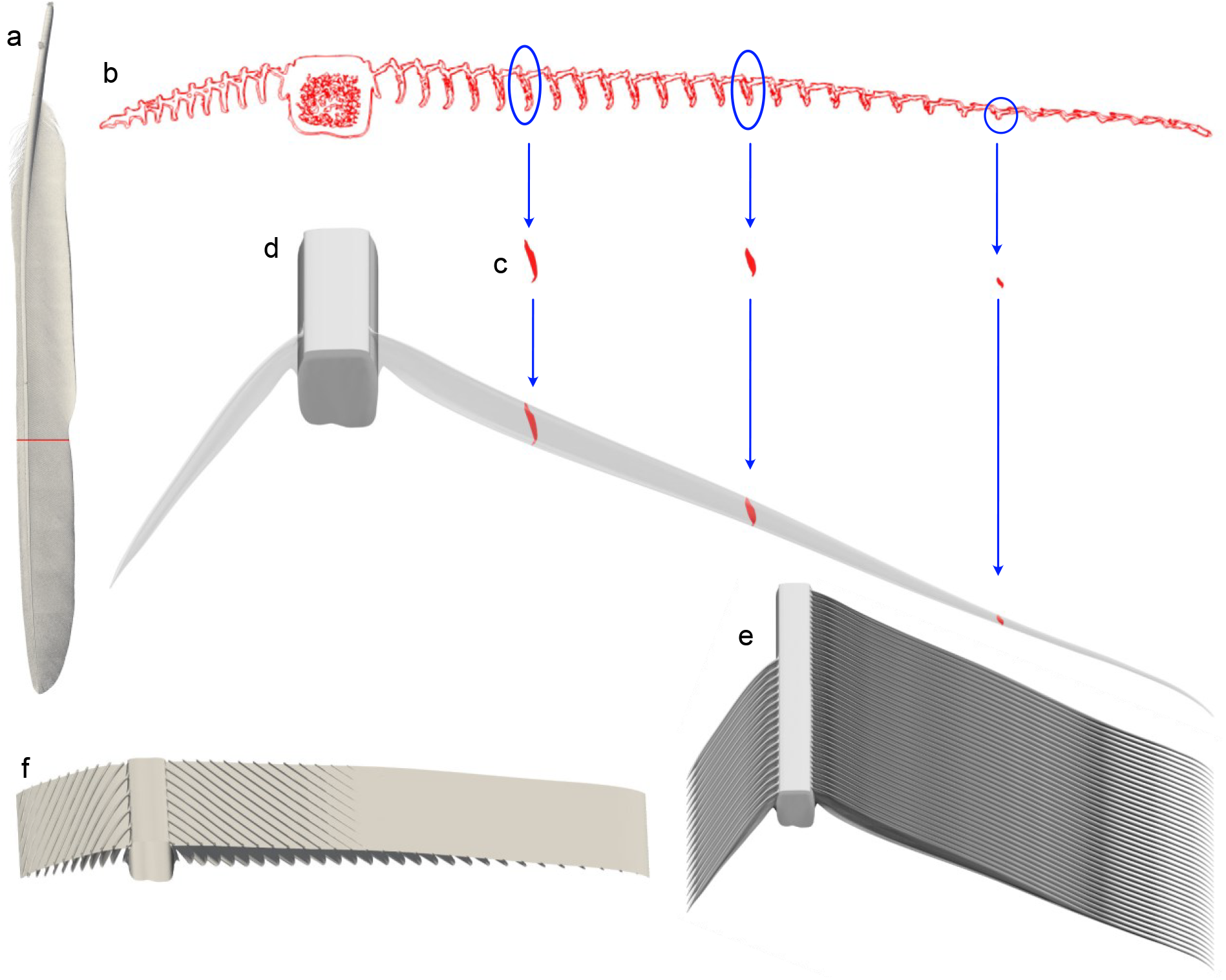
Process for creating the CAD model from a CT scan. *(a)* Showing the cross section (red mark) in the full feather CT scan, used as a template in the model. *(b)* The cross section of barbs (blue circles) is extracted and *(c, d)* translated to their 3D position along a single barb. A volume, creating a single barb, is then extruded connecting the different cross sections. *(e)* The 3D barb is copied and repeated, with the same spacing as the original scanned feather, to fill the entire width of the model span. The same process is used to create both trailing and leading vanes. *(f)* With the width of the model selected, a section of the feather is then cut and used for the modelling.

Size and speed are key components defining the behaviour of the air moving across the bird, their wings, and feathers. Birds range almost four orders of magnitude in size (from ~2.5 g to ~15 kg) and fly at speeds ranging from 0-20 m/s ^11–13^, resulting in different flow conditions which is captured by the Reynolds number (*Re*). Reynold number is a measure of the relative importance of inertial and viscous forces in the flow and based on an wings chord is calculated as:

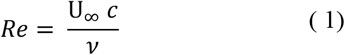

When categorizing bird aerodynamics, the size reference used are either focused on the entire bird ^12,14^, or even more often the wings ^15,16^ resulting in a *Re* of <100000-500000 ^12,13^. For the primary feathers, that function as individual wing, *Re* is much lower since the chord is smaller, and for the feather of interest here, in the order of 7000. The lower the *Re*, the more viscous (laminar) the flow behaves, meaning that the shear stresses in the flow are high enough to withstand disturbances ^17^. The flight mode also impact the flow conditions around the wings and feathers, and the two main flight modes, gliding and flapping flight ^12,13^ result in steady or cyclically variable flow respectively. Jackdaws have been observed to use both these flight modes and have been demonstrated to be competent gliders in the lab ^14^. In both these modes the flight feather used here form the leading edge in a slotted wing tip section (Figure 1) and work as a wing on its own.

Standard symmetrical aerofoils and plates at the range of *Re* relevant to feathers ^18–22^ display several fluid phenomena, including, but not limited to, laminar separation bubble ^18–20^, flow separation ^18–21^ and several modes of vortex shedding ^18,21,22^. There is also a strong three-dimensional aspect to many of the flow structures found around infinite wings at these low *Re*, as reflected in different results when comparing 2D and 3D flow measurements/simulations ^18,19^. Depending on the shape, such as thickness and camber, of the aerofoil or plate, the above mentioned phenomena occur at different angles of attack, which affect the lift and drag of the aerofoil ^23^. Currently, our understanding of when and why these flow phenomena occurs and how they are triggered is relatively limited. Even modelling and experimental results differ between studies of the same aerofoil under the same flow conditions ^18–20^, without any obvious explanation to why, suggesting results are sensitive to flow conditions and measurement methods. Interestingly, the profile shape of a feather (Figure 2b and Figure 3) does not resemble any of the manmade structures studied this far and it is not clear how the shaft and barbs affect the effective profile and flow around the feather that in turn affects the aerodynamic performance. This is especially interesting since the studies of aerofoils and plates in this low *Re* range indicate a sensitivity to the flow conditions ^24,25^.

**Figure 3.**
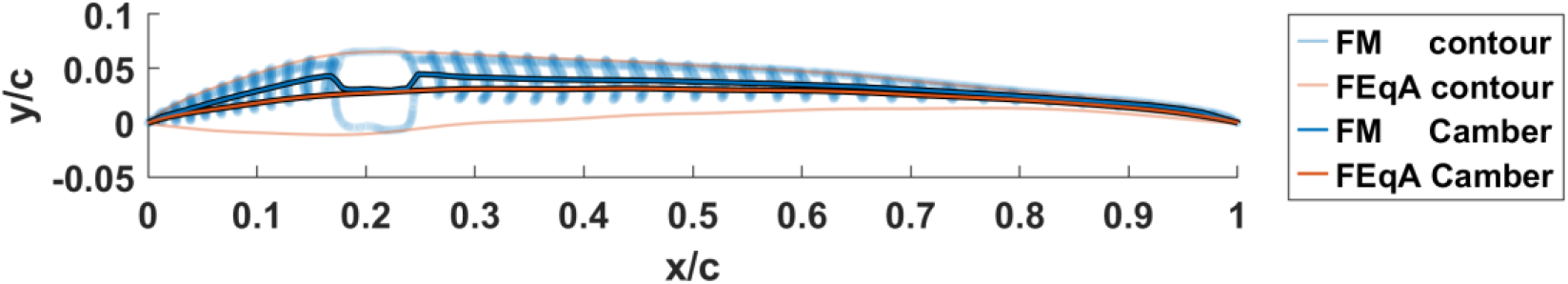
FM (Feather model) and FEqA (Feather equivalent aerofoil) profiles and camber. The transparent blue shows the FM with the dark blue line as the centre line. The transparent orange curve represents the FEqA surface with a dark orange centre line. The FEqA encapsulates the effective fluid surface of the FM at its zero-lift angle of attack.

Primary flight feathers have a strong three-dimensionality in their structures, where the shape and components, barbs especially (Figure 2), are laid out in such a way that 3D effects in the flow are likely to occur at any point of the feather which is exposed to free flow. In such cases, a 2D representation of a feather is impossible, and to the best of our knowledge, a detailed aerodynamic analysis of even a section of a feather has not previously been performed. Consequently, we do not know how feathers perform aerodynamically relative to standard foils, plates, and other flying animals (e.g. dragonflies). As a first step in understanding feather aerodynamics, we here performed a detailed computational fluid dynamic analysis on a model which represents a section of a jackdaw primary flight feather in gliding flight conditions with the goal to understand how the feather structures contribute to performance. We found that the protruding shaft and barbs above and below the barbule plane has a positive impact on the lift but a negative impact on the drag. The feather model shares several aerodynamic behaviours with standard aerofoils and plates, however, no single aerofoil that we compared with covers all the feather behaviours as the angle of attack changes. These behaviours include the shedding patterns, the chordwise movement of the centre of pressure and the consequent behaviour of the pitch torque, indicating a focus on flow stability rather than low drag in the feather section.

## 2 Method

### 2.1 Model creation

We based our feather model (FM) (Figure 2f) on a CT-scan of the ninth primary flight feather of a jackdaw (*Corvus monedula*) (Figure 2a). This feather forms the leading edge of the wing in the outer, slotted region of the wing (Figure 1). We performed the scan using a ZEISS Xradia XRM520 Submicron Imaging System (http://www.zeiss.de/) at the 4D Imaging Laboratory, Division of Solid Mechanics, Lund University with a voxel resolution of 10 µm. A high-resolution model was needed in the CFD simulations, to resolve the complex geometry, and due to the resulting computational time, we decided to only model a section of the feather. We used a cross section at 63% of the length, measured from the base, of the feather (Figure 2a), which is located where the feather is unsupported by its neighbours and therefore acts as an independent aerofoil. A CAD-model was constructed using SolidWorks2020 (Dassault systems Waltham, Ma, United States), so that the structures, shaft barbs and barbules, match the CT scan, at both leading and trailing vanes.

The shaft profile was extruded to the selected depth of the model. For the barbs, the relative positions of the cross-sectional profiles were translated along the spanwise direction to create a three-dimensional CAD model of a single leading and trailing barb, respectively (Figure 2c). The translation of each profile was determined to match the location of a single barb in the original feather scan. The trailing barb profiles are also translated vertically, to match the mean top position of the barbs, based on the average position of the CT scan and 3D surface scans of lower quality of four additional feathers. The additional scans were performed at the 3D-lab of the Department of Biology at Lund university using a 3D surface scanner (GOM ATOS I SO, GOM scan, Braunschweig, Germany). The single barb, on each side of the shaft, was subsequently copied and repeated to create the vanes (Figure 2e). The angle of the barbs at the front vane is 17° relative to the shaft and the barbs are spaced 0.80 mm apart (see supplementary material Figure S1). At the trailing vane the barbs have an angle of 29° relative to the shaft and a spacing of 0.50 mm, matching the original scan. The interlocking nature of the barbules from adjacent barbs to form the aerodynamic surface of the feather turned out to be too difficult to model and we instead opted to model the barbules as a solid plane, with 1.5×10^−3^c (0.0198 mm) thickness. The vertical position of this plane was fitted to the top barbule position on the barbs, which varies along the chord (see supplementary material Figure S2). On the trailing vane, the height of the barb above the barbule plane is continuously decreasing and due to resolution issues of the model, the top surface of the FM is modelled as a smooth surface for x/c > 0.5. The leading and trailing edges of the barbule plane were modelled as a pointed triangle. The spanwise length of the three-dimensional FM section used for the analysis is chosen so that it is continuously repeated (i.e. the barbs enter and exit the section along the span at the same chordwise location), to allow for cyclic boundary conditions. The final model had a chord of 13.2 mm and a width of 4 mm.

The initial analysis on the FM indicated areas with stationary recirculating flow, on both sides of the shaft and between the barbs on both the top and bottom side. To evaluate the effect of the surface properties of the feather we constructed a feather equivalent aerofoil (FEqA) (Figure 3), where these recirculation areas are included in the model so that the flow is effectively passing around the FEqA while maintaining the same behaviour as the FM. The top surface on the FEqA is based on the max vertical positions of the barbs above the barbule plane, the top of the shaft and the barbule plane at x/c > 0.5, creating a smooth top surface. The bottom surface is based on when the divergence of the Lamb vector ^26^, *Lamb* =ω × *U*, is 0. This condition indicated that the flow is irrotational ^26^, and the aim is to incapsulating all of the possible disturbances close to the feather surface and thereby creating a surface at the same location as the effective “fluid surface” is located. The FEqA bottom surface follows the iso-surface for the majority of the chord and deviates from it in front of the shaft and at the trailing edge to maintain a more aerofoil like shape. The FEqA is created for the flow condition at α=−2.59° (zero lift angle) of the original feather model. All in all, the FEqA have a smooth top and bottom surface, is slightly thicker than the feather shaft at its thickest point with the maximum thickness located in front of the shaft.

We determined the camber, the profile curvature, (Figure 3) by taking the difference between the top and bottom surface of the FM, defined by interpolation between the peaks of the barbs to get an even distribution of the surface every 5×10^−3^ x/c. FM has a camber of 4.47% and even though the top surface has the same curvature in the two models, the bottom surface is moved downwards for the FEqA, compared to the FM, and hence the camber line is also moved, resulting in a lower camber (3.15%).

### 2.2 Simulation and Mesh

#### 2.2.1 Domain and mesh settings

The total domain extended 5c above and below the centre of the shaft and 5c in front and 10 c behind the centre shaft (see supplementary material, Figure S3a)). We created the surface and volume mesh using Hypermesh 2021 (Altair, Troy MI, United States). We used a tetrahedral mesh, (see supplementary material, Figure S3 b-c)) with the FM surface mesh set to have an edge deviation of 0.15-0.38% of the chord and a growth rate of 1.05. The settings were used to ensure a resolution of at least 4 cells on the barbule plane between the barbs, in the streamwise direction. The height of the barbs on the top side was resolved with 5 cells and on the bottom side with 20 cells, closest to the shaft. Walls were placed at 0.75 c in front of, 1.25c behind, 0.5c above and 0.5c below the centre of the shaft to control volume mesh size near FM and FEqA, the walls themselves are not transferred to the simulation and are only used to build the mesh. The resolution at the surface of these walls was set to 3% of the chord. The size of the outer boundaries were set to 6% of the chord and the sides of the outer box were automatically set.

### 2.2.2 Numerical settings and boundary conditions

The computational fluid dynamic simulations were done with OpenFOAM8 (openfoam.org), solving the Navier-Stokes equations as a PISO implicit unsteady simulation, with no explicit turbulence modelling. Since a typical gliding speed for a jackdaw is around 8 m/s ^14^, we matched the chord-based *Re* (=7000). At this low *Re* the flow and boundary layer are laminar, but if the flow separates it is possible to have a turbulent wake, which makes it unsuitable to use RANS modelling. The boundary conditions on the side of the domain were cyclicAMI, which connects the two sides creating an infinitely wide domain to allow the flow to move in the spanwise direction along the barbs. The inlet, outlet, top, and bottom conditions are set to free stream velocity with the streamwise, and vertical velocity components set to match α and the FM and FEqA are modelled as walls. Given the distance between the model and the domain walls, these settings give similar results as setting the conditions to zero-velocity gradient. The temporal discretization time scheme was set to Euler scheme (i.e. 1^st^ order) for the first 0.5 seconds and then switched to backwards scheme (i.e. 2^nd^ order) and the convective term was discretized with a central order scheme (limitedLinearV0.1, due to simulation stability reasons), and the diffusion scheme was set to second order Gauss linear corrected. With these settings on time and space, turbulence should be captured when present. The free-stream velocity was set to 0.8 m/s and the time step was set to 0.1 ms, and the *Re* is matched by scaling the model by a factor of 10. We ran the simulations of FM at a subset of angles of attack ranging between −4.59°<α<20.41° and FEqA −2.59°<α<15.41° (avoiding the most extreme angles where the FM had problems generating stable mean values of C_l_, see below/supplemental information), with each angle requiring approximately 10000 CPU hours. For the most extreme angles of attack (α<−2.59° and α>12.41°) we did not manage to get stable means of the force coefficients over the sampling times used. This may have at least three different causes. First, we may not have run the model for long enough to stabilize the results, as a result of limited resources. Second, the means are taken over too short intervals to accommodate the low frequency oscillations that we see in the data. Third, the flow is truly chaotic and we may not expect the mean to stabilize over time ^27^. Since the third option exist, we decided to present the results in the figures to show trends in the behaviour of the FM, but note that the means are not accurate, and we caution against over-interpreting the results for extreme α.

#### 2.2.3 Mesh Sensitivity

Mesh sensitivity was determined (see supplementary material Figure S4), using three surface mesh resolutions, one with half the surface resolution we finally used, one with the final mesh resolution and one with twice the surface resolution of the final mesh. Simulations were performed for α=0.41°, α=4.41° and α=5.41°, and the C_l_ and C_d_ were calculated as the averages of the last second of simulation. Across the three angles tested the results (see supplementary material Figure S4) of the coarse mesh shows the most difference from the other two and the C_l_ deviation between the final mesh to the high resolution mesh was less than 1 percent for α=0.41° and α=5.41° and 3% for α=4.41°, the C_d_ deviation was less 2% for α=0.41° and α=4.41 and less 3% for α=5.41°. Due to the geometric complexity of the model increasing the resolution of the model is not necessarily expected to result in asymptotically increasing or decreasing lift and drag. Instead increasing resolution is expected to reduce the variation in the results. In our case the difference between the final mesh and the high-resolution mesh are within a few percent, even at the angles of attack where flow characteristics are most sensitive to resolution. Given the computational resources available we therefore decided to use the intermediate mesh resolution for the simulations. We also conducted a secondary mesh study in combination with the numerical settings (see below). Unlike the study of the FM mesh resolution that study focused on the resolution of the flow in the domain, which showed a corresponding response to the reference case.

#### 2.2.4 Numerical validation

We performed a validation study of the simulation settings (see supplementary material Figure S5) using an Eppler E387 at Re=10000. These simulations were performed in OpenFOAM 10. The base case has the following properties: The chord length is c=1 and the leading edge of the aerofoil is located 6c from the inlet. From inlet to outlet the length is 16c and in the cross-stream direction 10c. The width of the span is 0.4c. The mesh is generated using snappyHexMesh and the aerofoil is surrounded by 2 refinement boxes with cell size dx/c=2.5 ×10^−2^ and dx/c=1.25 ×10^−2^ respectively, and closest to the surface there are 6 layers with a resolution of dx/c=1.56×10^−3^. The convective terms are discretized using the upwind scheme (i.e. 2^nd^ order), and the temporal discretization is done using the backward scheme. Angles of attack in the range from −6 to 18 degrees were considered. In addition to the base case, we studied the influence of mesh resolution, discretization scheme and domain size. The mesh resolution was studied with one case with dx/c=7.81×10^−4^ (half base case size) closest to the surface and one case with a prismatic refinement on the original mesh. Additionally, another convective scheme has been tested, with a 2^nd^ order central scheme (limitedLinearV 0.1, same as FM settings), as well as two cases where the domain size was considered, one with doubling the size in length and height directions and one with a doubling in the span-wise direction. In all cases, the simulated time was 144 c/U. The flow was allowed to develop for 16 time units and C_l_ and C_d_ were then average over the following 128 time units. The time step size was either 8×10^−4^ c/U or 4×10^−4^ c/U depending on the mesh resolution and the numerical validation (see supplementary material Figure S5) study of mesh resolution, numerical discretization, and domain size show a robust response to the changes made, with little deviation to the base case. For validation, experimental data of E387 ^24^ was selected, however, afterwords, we noted that this particular data stands out in performance relative to other airfoils (i.e. Cl & Cd deviates by a factor ~1.2). Simulations on additional profiles, E61 and NACA4403 (see supplementary Figure S6), and computational results of the same profile ^28^ also reinforces this notion. The numerical validation shows our model to be stable and alternative simulations of the same profile ^20,29^ support our conclusion, suggesting that additional experimental studies of the E387 profile at low Reynolds number would be desirable.

### 2.3 Extraction of results

If nothing else is stated, the results presented and discussed are the time averaged results of the last 1.5 seconds of the simulations. If instantaneous results are presented with no time indication the data is collected from the last time step of the simulation of the 1.5 s collected for the time average data. Data presented as a 2D section was extracted at a span position of 40 percent of the model. Additionally, velocity along lines perpendicular to the chord were also extracted at this position, where the lines are 0.23c high, cantered around the midpoint of the shaft and are at equidistant positions along the chord with a spacing of 0.104c and starts at leading edge of the FM. Data was further processed in Paraview (paraview.org) or Matlab r2022a (Mathworks, Natick MA, USA).

Force coefficients comparison with other studies have been enabled by using plotdigitizer (plotdigitizer.com) and extracting values by uploading images of graphs into the web-based app and manually selecting data points.

#### 2.3.1 Force and pressure coefficients

Force coefficients (C_l_ and C_d_) were calculated according to Eq. (2) and Eq. (3), where the lift direction is set perpendicular to and the drag direction is set parallel to the free-stream flow.

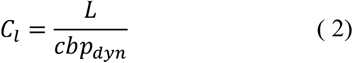

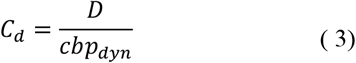

We estimated force coefficients as the mean values of the last 1.5 seconds of the force coefficient monitors, created by OpenFOAM8. To estimate the validity of the mean values the average of the lift has also been calculated for shorter time periods (0.1 – 1.4 seconds) (See supplementary material Figure S7b). The difference between the mean value with an evaluation time of 1.4 and 1.5 seconds lies between 0-2.5% for −2.59°≤α≤12.41° and between 2,5-17.5% for α<−2.59° and α>12.41°. The dynamic pressure is defined as to *p*_*dyn*_ =0.5ρ*U*.

The pressure coefficient was calculated according to Eq. (4).

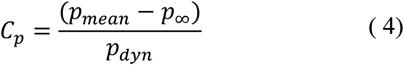

#### 2.3.2 Torque and centre of pressure

We determined the torque around the centre point in the shaft, Eq. (5), from the distribution of the aerodynamic force over the entire surface of the model. The centre of pressure was then estimated by dividing the torque by the normal force to the chord and the calculated distance was placed along a horizontal line running through the centre of the shaft. Using (5) the pitch torque coefficient (C_M_) has been calculated as described in Eq. (6).

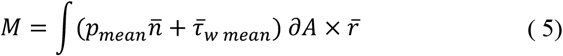

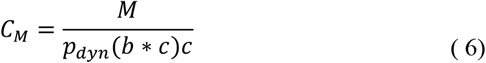

## 3 Results

### 3.1 Feather model (FM)

#### 3.1.1 Feather flow characteristics – top side

At the leading vane, the air flows above the barbs, but stays attached to the vane for α<12.41°. The flow between the barbs is directed at an angle to the freestream along the barb, flowing towards the root of the barb for α<12.41° (Figure 4). For the higher angles, α≥12.41°, the flow separates at the leading edge of the FM (see supplementary material Figure S8), resulting in a negative chord-wise (forwards directed) flow. The flow over the shaft reflects the situation in front of the shaft and stays attached when the flow is attached at the leading vane and detached when separated at the leading vane.

**Figure 4.**
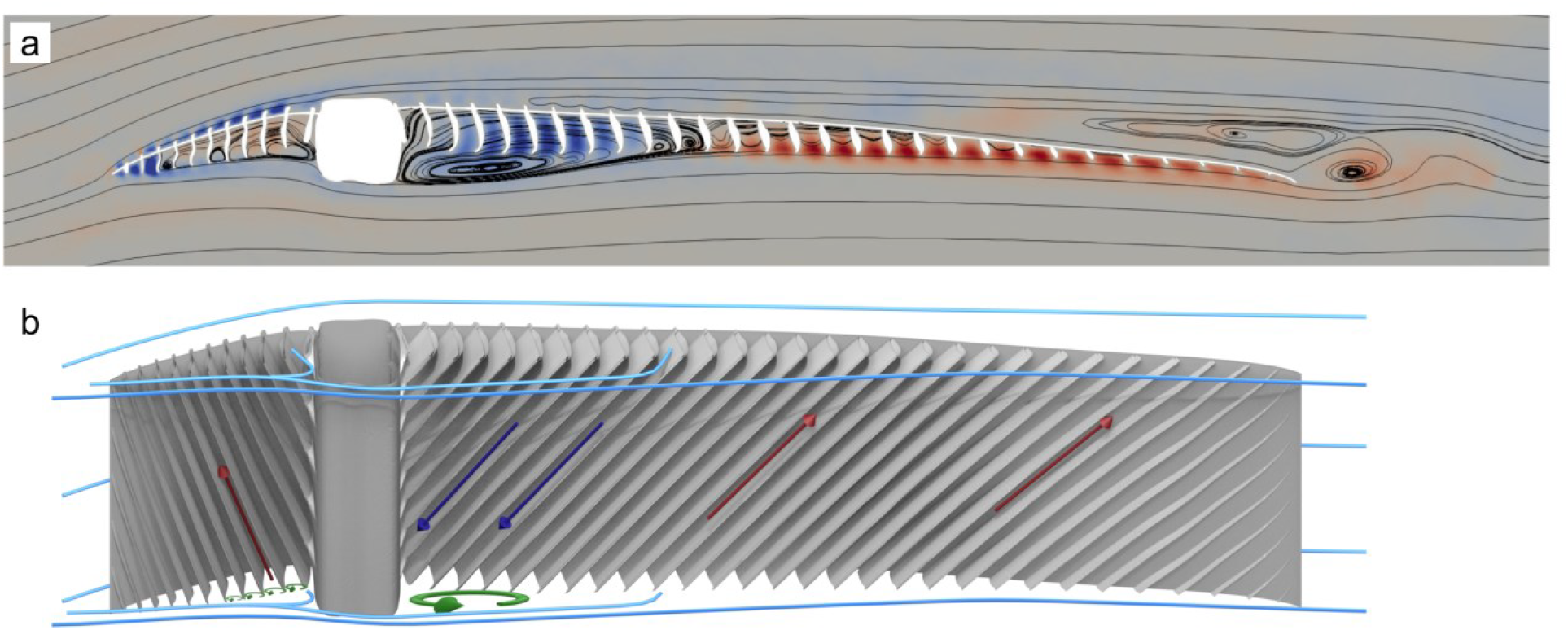
Flow directions around the FM at α=5.41°. *(a)* The streamlines overlay the spanwise flow, with positive values (out of the page) in red and negative values (into the page) in blue in a plane located at the centre of the model. On the leading vane, positive values indicate flow away from the shaft, while behind the shaft, on the trailing vane, negative values show flow towards the shaft. *(b)* Schematic image of the flow in *(a)* viewing the model obliquely from below. Blue lines show streamlines flowing from the left, red and blue arrows show the flow direction of individual flow structures, same as in (a), and green circle arrows show recirculating flow (see main text).

Over the trailing vane the flow is attached for −5.41°≤α≤0.41°, and at α=3.41° the flow starts to detach from the trailing edge. The size of the detached zone increases slightly at α=4.41° and at α=5.41° the entire trailing vane has negative chord-wise flow close to the barbule plane. Starting at α=3.41°, the flow between the barbs rotates streamwise around an axis parallel with the barbs, with the rotation centre at the same height as the top of the shaft (above the barbule plane). At α>5.41°, this rotational flow is no longer visible due to flow separation. The separated flow is characterized by no longer being attached to the surface and forming a recirculation region. Inside the wake the pressure and velocity are different than the surrounding flow decreasing lift and increasing drag.

The pressure distributions for α<10.41°, show a decreasing pressure from the leading edge to the shaft where the minimum is located (Figure 5a, b). The curve is jagged, showing variation in the pressure, due to the impact of the barbs. The pressure then increases towards the trailing edge. For α>10.41°, the location of the minimum pressure moves from the shaft to the leading edge of the FM, the pressure then increases over the leading vane until the shaft. Behind the shaft, the pressure distribution is level, with a small increase at the trailing edge, for α=10.41°. At α=15.41° we see a slightly different behaviour behind the shaft, where the pressure decreases at ~x/c=0.55 and then increases only to decrease at the trailing edge.

**Figure 5.**
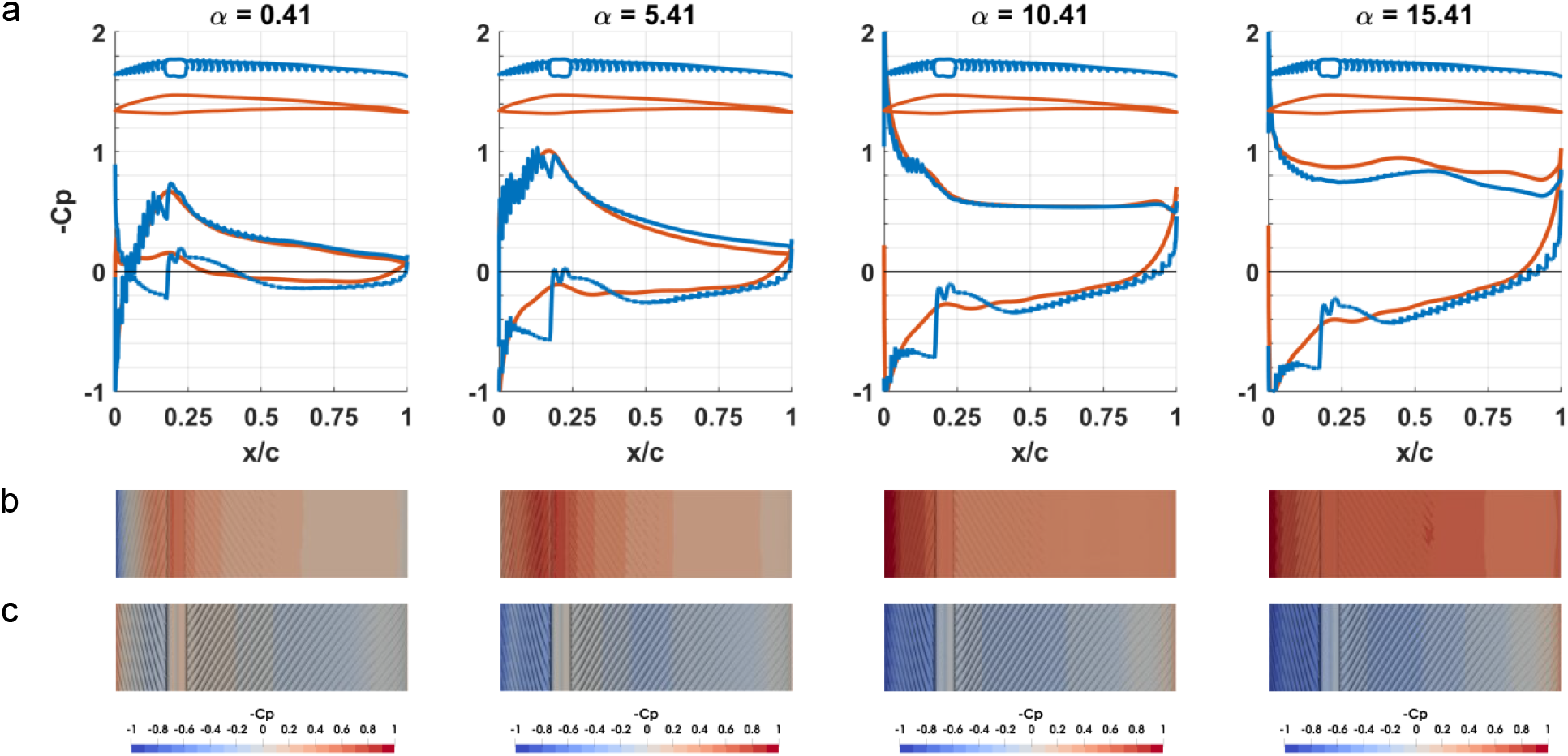
Pressure coefficients (C_p_) for FM (blue) and FEqA (orange) at α = 0.41°, 5.41°, 10.41° and 15.41°. *(a)* C_p_ for FM and FEqA. The outline of each cross section is shown above the C_p_-curves. *(b)* C_p_ plotted on the top surface of the FM. *(c)* C_p_ plotted on the bottom surface of the FM.

### 3.1.2 Feather model flow characteristics – bottoms side

The flow below the FM is more complex than on the top side. For α≥0.41° the protruding shaft creates a high-pressure zone in front of it and a low-pressure zone behind it (Figure 5a, c). The pressure distribution varies along the chord, with a high pressure at the leading edge followed by a pressure drop to a local minimum, only to increase again in front of the shaft. Over the shaft and directly behind it, the pressure is relatively low and then increases to create a local maximum at the trailing vane (at ~x/c=0.5) and then decreases towards the trailing edge (Figure 5a). As the angle of attack increases, the position of the local minima and maxima moves towards the shaft.

Since the distance from the edge of the barb to the barbule plane is longer on the bottom side than on the top side of the FM, rotational flow structures form between the barbs on the bottom side of the leading vane, for α>0.41°. The rotational structures are driven by the air flow beneath the barbs, but do not fill the entire space between the barbs (Figure 4a and see supplementary material, Figure S9), and instead form closer to the openings than to the barbule plane. As the flow continues and reaches the shaft, a stagnation point is created at ~1/4 of the shaft height, which does not move for α>0.41°. At the stagnation point the air is divided and part of the air flow is directed towards the barbs and barbule plane. As the flow reaches the barbs and enters the space between the barbs it is redirected to align with the barbs and flow away from the shaft, above the rotational structures, towards the local pressure minimum close to the leading edge of the vane (Figure 4). The rest of the flow is directed below the shaft and continues towards the trailing vane. The flow separates from the shaft and in the low pressure behind the shaft a single rotational structure fills the volume behind the shaft (Figure 4). Behind the rotational centre and up to the local pressure maximum at the trailing vane, part of the flow is directed upward towards the barbs and barbule plane, forcing the flow to move in between the barbs. The direction of the flow between the barbs is dictated by the local pressure maxima. Between the shaft and the local maximum, the flow is directed towards the shaft and downstream the local maximum the flow is directed away from the shaft (Figure 4).

#### 3.1.3 Force coefficients

Based on the response of C_l_, see Figure 6a), α can be divided into five different groups (see supplementary material Figure S7). The first group, with an α<−2.59° (below zero lift), is characterized by large amplitude temporal fluctuations in C_l_ due to flow separations on the bottom side of the FM. The second group, containing −2.59°≤α<5.41°, is characterized by low amplitude fluctuations in the force coefficients due to lack of any type of shedding. The third group, containing 5.41°≤α≤7.41° has clear and stable amplitude fluctuations in C_l_, resulting from a von Karman type of shedding at the trailing edge. The fourth group, containing 10.41°≤α≤12.41° also has clear, stable high frequency oscillations, but also includes a signal with lower frequency, superimposed on the “stable signal”. For α<12.41° we see unstable, potentially chaotic, oscillations in C_l_. Given that there is a possibility that the results represent the real circumstances, we decided to include them, but caution against over-interpretation of the averages.

**Figure 6.**
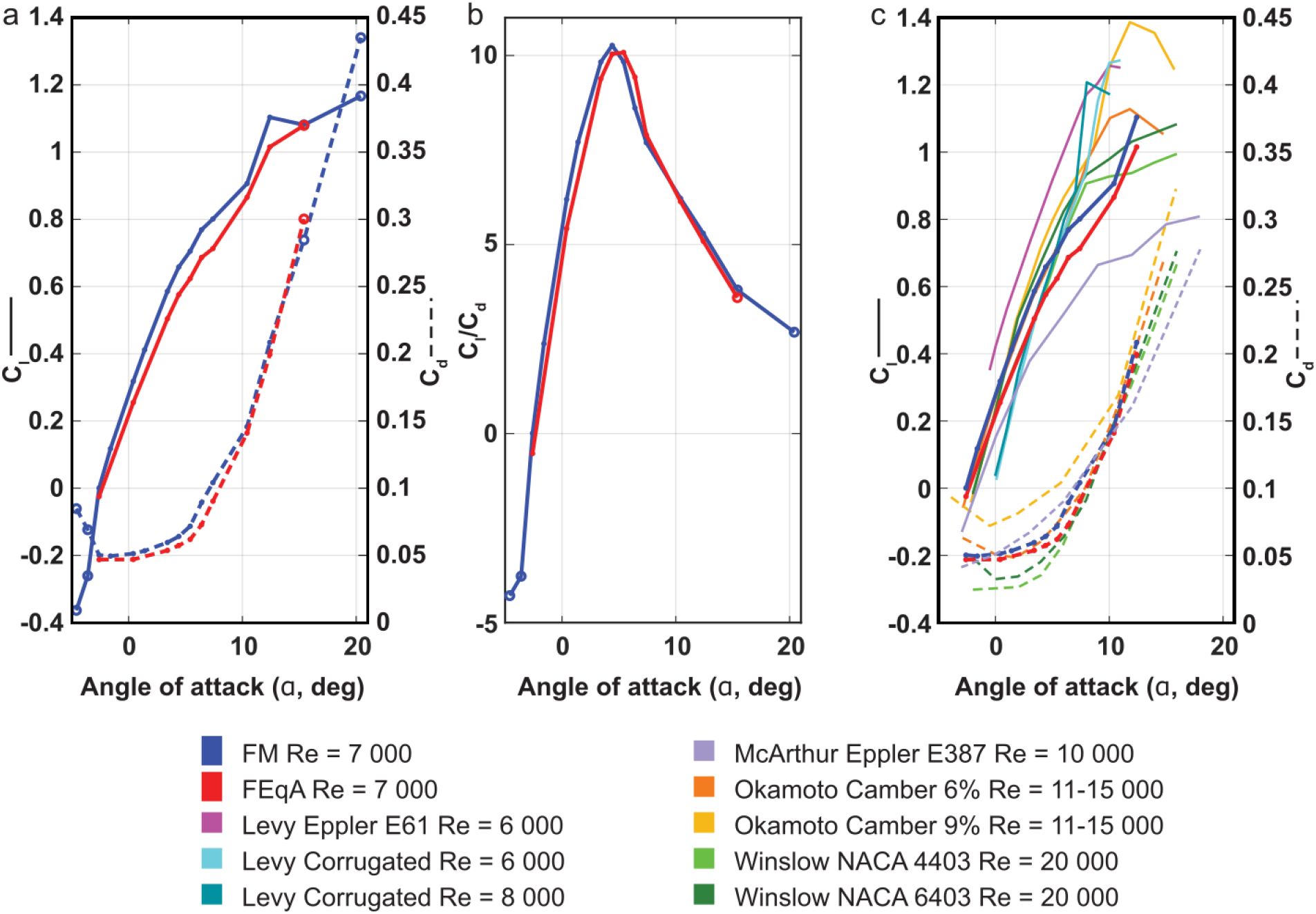
Performance of feather model (FM), equivalent aerofoil (FEqA) and other aerofoils which versus angle of attack. *(a)* Average C_l_ and C_d_ for the FM and the FEqA. *(b)* Glide ratio, C_l_/C_d_ for the FM and FEqA models showing similar behaviour but with slightly better performance for the FM. *(c)* Average C_l_ and C_d_ for the FM, as well as the FEqA in comparison to literature results from studies of aerofoils and cambered plates at relevant *Re* (6 000 – 20 000). References of experimental data from Okamoto ^23^ and McArthur ^24^ while simulation data are from Winslow ^20^ and Levy ^29^. Open circles in *a* and *b* mark data points where the change in mean data differs more than 2.5% between an average of 1.5 and 1.4 second (See supplementary material Figure S7). These points have been left out of the comparison with other data in *c*.

C_l_ becomes positive at α=−2.59° and then continues to increase with similar characteristics until α=10.41° at which point it increases rapidly before levelling off as α increases further (Figure 6a). C_d_ is slightly above minimum for the lowest angles of attack tested, decreases to a minimum at α=−2.59°, and then starts to increase as the angle of attack increases further (Figure 6a). The glide ratio (C_l_/C_d_), shown in Figure 6b), increases until α=4.41° where it peaks and then decreases rapidly until α=7.41° and subsequently tapers off.

#### 3.1.4 Shedding

As indicated in the C_l_ results, the flow separates from the FM for the lowest angles of attack α<−2.59°. However, for −2.59°≤α<5.41° the flow stays mostly attached to the FM and separation occurs only at the trailing edge. At α=5.41°, we start seeing von Karman-like shedding at the trailing edge, which is also indicated in the force coefficients, where the values start oscillating (Figure 7 and supplementary material Figure S7a). It is also at this α where the separation point moves from the trailing edge to the shaft (see supplementary material Figure S8). As the angle of attack increases further, the shedding associated with the bottom side of the FM continues to release from the trailing edge, but on the top side the shedding jumps forward towards the shaft and starts to release structures at about half chord. As the angle of attack increases, the shedding moves further forwards, although the flow separation point has not moved (see supplementary material Figure S8), and the flow still separates at the trailing point of the shaft at a=10.41° (Figure 7). At α=15.41° the flow separation point on the top side, has moved to the leading edge of the FM and the shedding start in front of the shaft (Figure 7). When large scale shedding occurs at the shaft/mid chord, smaller vortex structures can be seen moving along the top of the rear vane. These structures are three dimensional and rotate in the opposite direction to the larger scale structures found above them (Figure 7 just in front of trailing edge).

**Figure 7.**
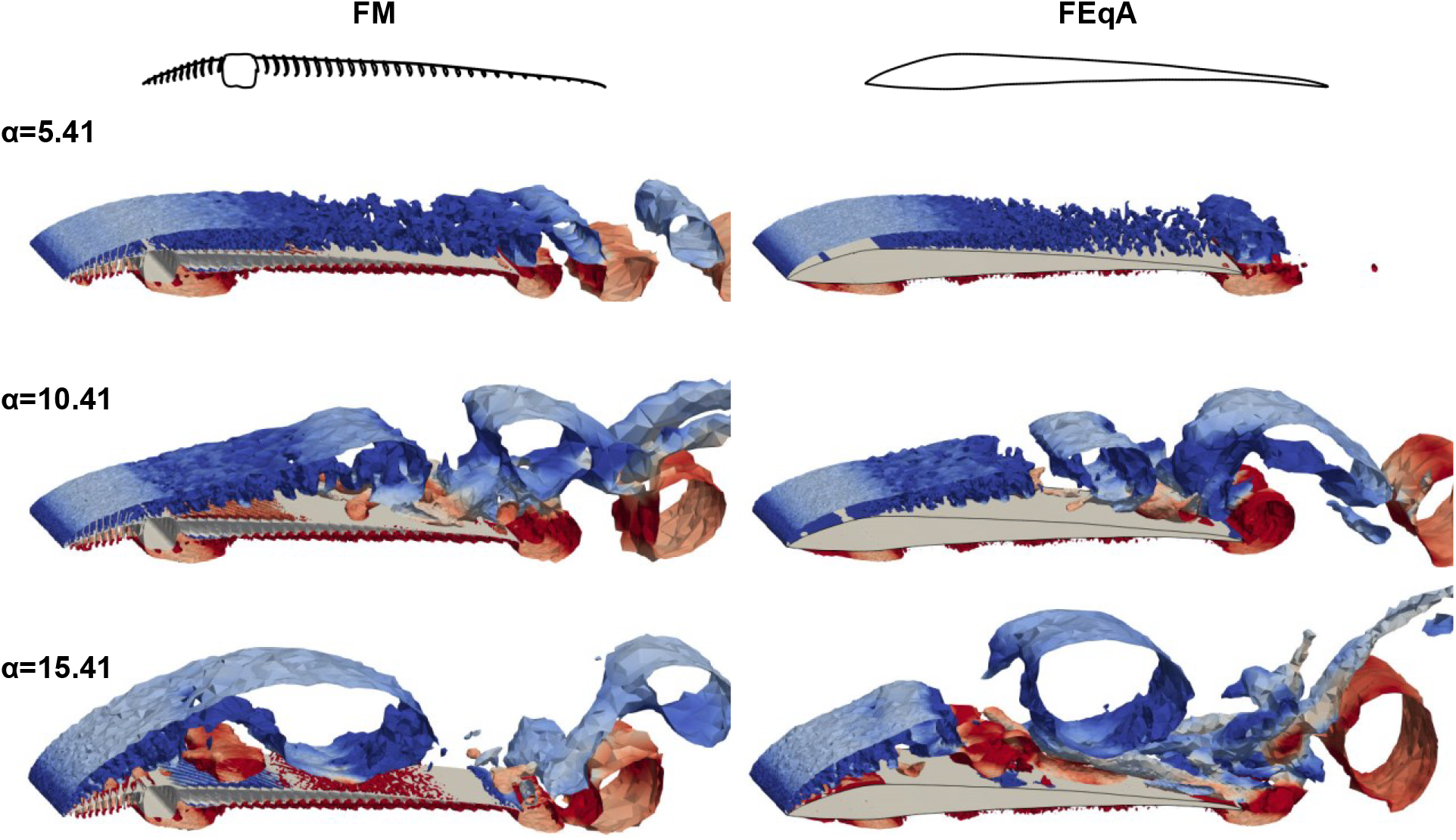
Instantaneous vortex structures shed from the FM and FEqA illustrated as surfaces of Q-criterion at level 50 for α = 5.41°, 10.41° and 15.41°. The vortices are coloured by the spanwise vorticity, clockwise as blue and counterclockwise as red. Animated results for FM at α=7.41° see supplementary material Movie M1. Note that the panels do not represent the same phase of the vortex shedding.

#### 3.1.5 Pitch torque around shaft

The C_M_ of the FM, calculated relative to the centre of the shaft (Figure 8a), is negative (pitch down rotation) for all angles evaluated. The curve has two distinct slopes (Figure 8b), one slope for −2.59°<α≤5.41° and a steeper slope for 5.41°<α<15.41°. The difference in slopes can be attributed to the movement of the centre of pressure (Figure 8 c, d). For large angles of attack, α>5.41°, the centre of pressure moves towards the leading edge as the angle is reduced, but as the angle of attack continues to reduce below α≤5.41° the centre of pressure instead moves toward the trailing edge. Note that for 5.41°<α<10.41° there are inconsistencies to this pattern where the centre of pressure does not strictly follow a straight pattern towards the trailing edge (Figure 8 e).

**Figure 8.**
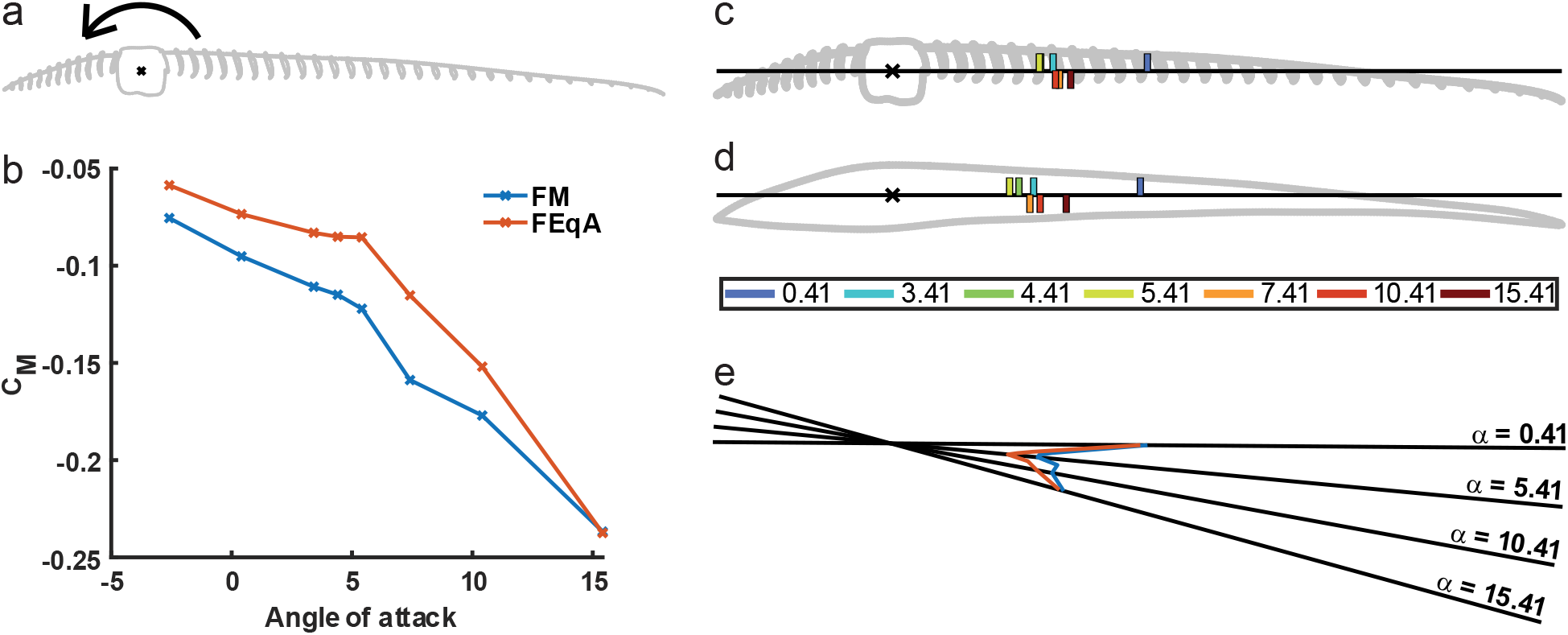
Torque around the centre of shaft and location of centre of pressure. *(a)* Rotation centre and pitch down direction of the feather. *(b)* C_M_ around shaft centre for FM (blue) and FEqA (orange). *(c)* Location of centre of pressure for FM, indicated by coloured bars. The numbers indicate the integer value of α. (e.g. 0 represents α 0.41) Locations of CoP α≤5.41° above the line and α>5.41° below the line. *(d)* Location of centre of pressure for FEqA, indicated by coloured bars. Locations of CoP α≤5.41° above the line and α>5.41° below the line. *(e)* The location of centre of pressure plotted on a rotated chord line (with rotation centre at shaft centre) to match relative position to shaft and α, FM (blue), FEqA (orange).

### 3.2 Equivalent aerofoil (FEqA)

#### 3.2.1 Flow around FEqA

The flow pattern above the FEqA is similar to the flow above the FM. The velocity profiles (Figure 9) have the same characteristics, e.g. shape of the profiles, and on the front vane they are overlapping. However, on the bottom side there are some dissimilarities, with the shaft in the FM having a large impact on the flow. The continuous surface of the FEqA allows the flow to follow its path more easily than for the FM, allowing the velocity in the boundary layer to build up to the ambient velocity more quickly than for the FM.

**Figure 9.**
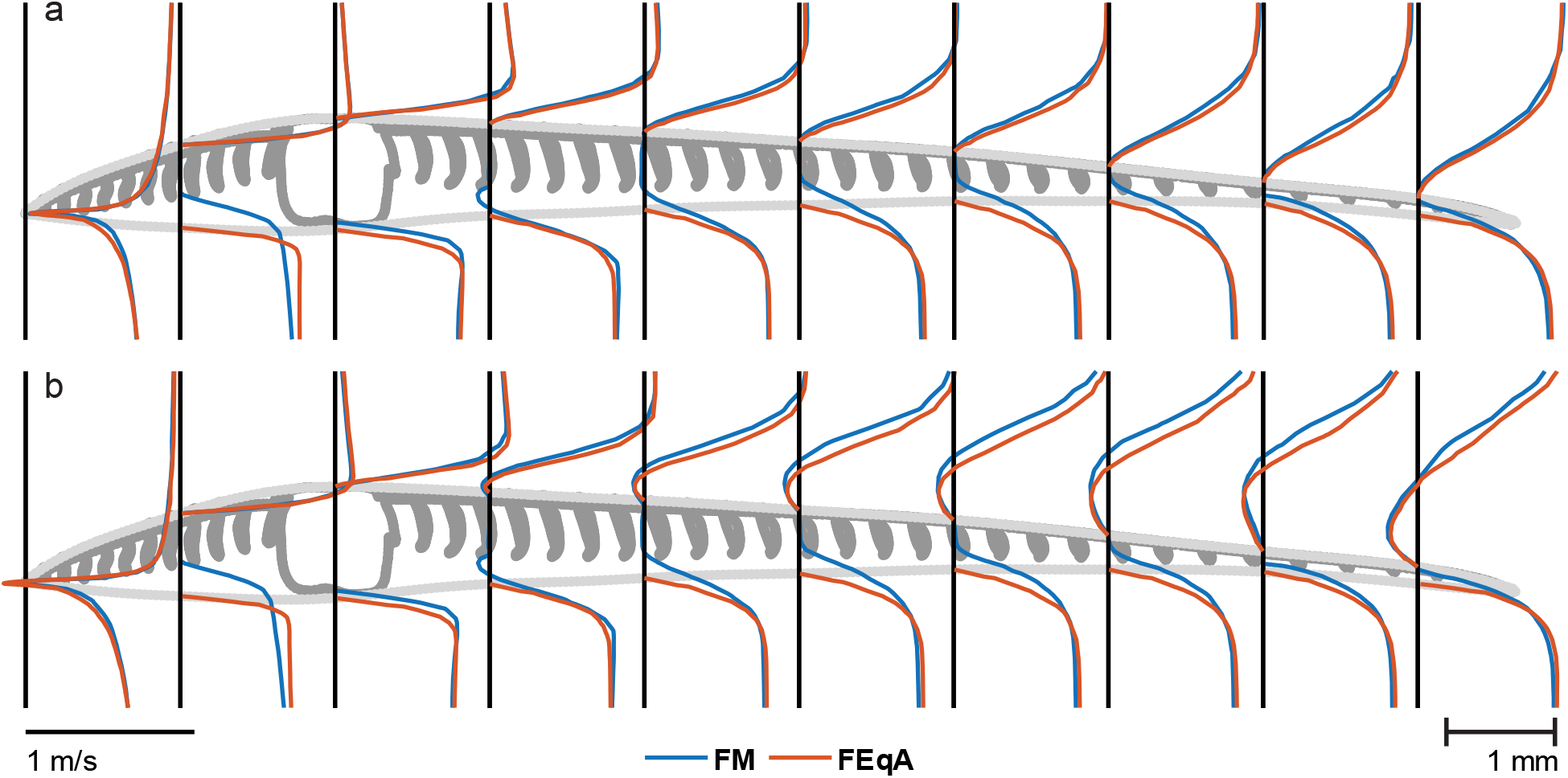
Velocity profiles for streamwise velocity component for FM (blue) and FEqA (red) at α=4.41° *(a)*, and α=7.41° *(b)*. The velocity is zero at the surface of each profile, where the vertical black lines meet the models. Velocities are scaled according to the line in the lower left corner of the figure.

Unlike the FM, the separation point on the trailing vane on the top side of the FEqA moves towards the shaft in increments as the α increases (see supplementary material Figure S8). At α=4.41°, a separation occurs at the trailing edge, while at α=5.41°, the separation point has moved to ~2/3 of the trailing vane, and at α=6.41°, the separation point is even closer to the position of the shaft in the FM. For 7.41°≤α≤10.41°, the separation occurs in the vicinity of the shaft position and for α=15.41°, the flow separates at the leading edge.

#### 3.2.2 Pressure

The difference in flow between the FM and FEqA models is also reflected in the pressure distribution. The C_p_ curves (Figure 5a) on the top side are similar between the two models, where the main difference is found on the leading vane. Here, the barbs cause the pressure to build in front of them and decrease behind them, creating a jagged pressure curve (Figure 5a). The removal of the barbs in the FEqA smooths the pressure curve, but the curves of the two models have the same overall characteristics.

On the bottom side there are significant differences between the FEqA and the FM, especially in front of the shaft, where a significantly higher C_p_ can be seen for the FM (Figure 5a). At the trailing vane there are also positive effects for the FM with higher C_p_ values than for the FEqA. The only position along the chord where the FEqA shows better performance than the FM is behind the shaft, where it lacks the low pressure found in the FM.

#### 3.2.3 Shedding and lift and drag

C_l_ of FEqA (Figure6a) shows the same grouping behaviour with α as the FM (see above, and supplementary material Figure S7 and Figure S10). One difference is that the FEqA has not been investigated for the lowest α, i.e. α<−2.59° so the first group is not detected. Another difference is that the start of the von Karman shedding has moved from α=5.41° to α=6.41°, coinciding with when the separation point moves to the shaft region for both models. The C_l_ of the FEqA is lower than for the FM, as is the C_d_ for all angles but α>12.41° (Figure 6 a), where C_d_ is higher than for the FM, e.g. at α= 4.41, Cl was 12.5% lower and Cd 10.5% lower than for the FM. C_l_/C_d_ of the FEqA is lower than or the FM due to the relatively lower C_l_ (Figure 6 b). However, the shape of the peak is wider than for the FM, due to the better C_d_ values for 5.41°≤α≤7.41° associated with a delayed start of the von Karman shedding until α=6.41° for the FEqA. The overview of the instantaneous vortex pattern (Figure 7) shows similar patterns between FM and FEqA. Signs of no shedding (FEqA) can be seen at the trailing edge of α =5.41°, otherwise the similarities of the separation point make the patterns similar.

#### 3.2.4 Centre of pressure and torque

The pitch torque across angles of attack (Figure 8 b) for the FEqA follows the same pattern as for the FM with the curve following two distinct slopes, one from −2.59°≤α≤5.41°, and a different for 7.41°≤α≤15.41°. The main difference between the two models are the values of the slopes, where the FEqA has lower values than the FM. Additionally, the curve is smoother for the FEqA than for the FM in the range 5.41°≤α≤15.41°. The FEqA also follows the same movement pattern for the centre of pressure as seen for the FM (Figure 8 e). The centre of pressure moves from the trailing edge towards the shaft when α is decreasing for 5.41°≤α≤15.41° and from the shaft towards the trailing edge as the α continues to decrease for 0.41°≤α≤5.41°. The inconsistent pattern seen for the FM, at 5.41°≤α<10.41°, is less noticeable for the FEqA.

## 4 Discussion

Feathers have been subjected to a complex selection pressure in relation to the various flight related functions during evolution ^10^. A primary flight feather, in a split wing tip, not only has to generate aerodynamic forces efficiently and effectively as a standalone unit during both gliding and flapping flight, but it should also transfer these forces to the wing without breaking and preferably use as little material as possible to keep the weight of the wing and bird low. Given these demands on the feather, we may not expect that aerodynamic efficiency (e.g. C_l_/C_d_) is necessarily optimal in feathers, but instead the other demands, such as structural stability, may supersede in importance. Our study provides a first indication of this trade off by determining the aerodynamic performance of a section of a feather and the relevance of the microstructures of feathers in gliding flight conditions. We have found that the FM is effective at producing lift, with a performance comparable to aerofoils and plates with larger camber and higher Re ^20,23,24,29^ as well as to our FEqA (Figure 6c, d). The shaft and barbs do not have an apparent negative impact on C_l_, but has a negative effect on C_d_, compared to the smooth FEqA (Figure 6a). This indicates that other functionality, e.g. structural demands of the shaft may be more important than the cost associated with a lower aerodynamic efficiency, i.e. a higher C_d_ and consequently a lower C_l_/C_d_. Furthermore, the FM and dragonfly wing show a low, relatively constant variation in C_l_ generation, at low α over time, while the variation systematically increases in the technical aerofoils, (see supplementary material, Figure S7a and Figure S11), indicating that the FM and a dragonfly wing^29^ behave more similar at low α, than compared to aerofoils and plates ^18,20,24^. This similarity may, given the disparate origins of birds and insects, but similar *Re*, suggest a potentially common pattern in biological systems; to prioritize low fluctuation and predictable forces. By extension, if true, this indicates a relative evolutionary importance of flow control in animal flight, which merits further investigation. Given that it is well known that in the *Re* range of animal flight the aerodynamic behaviour of aerofoils is very sensitive to the flow conditions ^20,29^, the potential for feather structures to mitigate this sensitivity should provide inspiration for future studies.

Although the FM generates C_l_ similar to FEqA (lacking microstructures and more similar to traditional aerofoils), the flow around the FM is different. On the bottom side, the shaft of the FM greatly impacts the flow (Figure 5 and 7) and prevents air from following the bottom of the barbs, creating a high-pressure zone in front of the shaft and a low-pressure zone behind the shaft (Figure 5), indicative of added drag. On the top side of the FM, the surface is not smooth, with the barbs and shaft protruding above the barbule plane (Figs 2 and 4). Accordingly, when removing the structures disturbing the flow in the FEqA, we find an improved C_d_ relative to the FM (Figure 6a). On the other hand, there is instead a relative reduction of the C_l_ (Figure 6a), resulting in lower C_l_/C_d_ for the FEqA than for the FM (Figure 6b), since the improvements in C_d_ cannot compensate for worse C_l_. The difference in performance between FM and FEqA may, however, also be related to a lowered camber for the FEqA compared to the FM, since a lower camber is associated with lower lift ^23^. At the same time, the shaft and barbs on the top side of the FM fixes the separation point (see supplementary material Figure S8) to the shaft region for low α, increasing the wake size (Figure 9) and therefore also C_d_, thus suggesting a negative effect of the surface microstructures on the C_d_ of the FM, but also a predictability in the separation point location. The difference in separation point location is also detectable in the instantaneous snapshots of the flow (Figure 7), where the shedding patterns are similar when the separation points align. When investigating the vortex shedding over the FM’s top side, unsteady structures and vortex shedding patterns are similar to that of aerofoils ^18,22^, with the exception of the laminar separation bubble ^18–20^ which we did not detect on the FM (see further discussion in supplementary material). Overall, this indicates that, the shaft has a relatively small negative effect on the performance of the FM Comparing C_l_ of the FM to engineered aerofoils may be precarious given the low Re ^25^, which is known to generate differences between simulated data and experimental data, as well as differences between different experimental testing sites for the same foil and Re ^24,25^. That said, we note that for flat plates (at Re 11000-15000) to have C_l_ values comparable to those of our FM, a higher camber than in FM and FEqA is required ^23^ and aerofoils need to operate at a higher Reynolds number (Re = 20000 than used in our study. However, these aerofoils are thicker than the FM, which gives lower C_l_ at low Re ^23^. Our validation test, of the simulation settings and the domain size, (see supplementary material Figure S6) concludes that the model setup is stable but possibly overpredicting C_l_. At the same time, the same model setup for NACA4403 and E61 (see supplementary material Figure S6) gives results in range with Levy and Winslow ^20,29^ presented in Figure 6, which suggests validity of the method. The possible lower aerodynamic performance of the FM at higher α than modelled, would suggest that the aerodynamic performance is not as evolutionary important as suspected or the contribution to the overall flight performance is not as important. However, if the FM can provide similar C_l_ as engineered aerofoil designs (Figure 6), after 150 Mya of evolution has shaped the morphology of the FM, then it leaves room for other potential benefits, such as structural stability (larger diameter shaft) and weight minimization. Low weight is also beneficial when considering other flying modes (e.g. flapping flight and manoeuvres), which increase the effects of inertia. Flapping is also likely to increase peak forces and thus the structural demands, further supporting the notion that structural properties has been a strong driving force in the evolution of feather morphology.

In addition to the lift and drag, the pitch torque around the shaft is affected by the changes in α (Figure 8). The torque around the centre of the shaft is always in a pitch down direction for the tested α, since the centre of pressure is always behind the shaft for α>-2.59° (zero lift). The combination of the torque arm and development of C_l_, results in two distinct slopes in the C_M_ curve vs. α (Fig 8b), with a stronger pitch down torque for α above peak C_l_/C_d_ than below. However, there are a few points not following this pattern (Figure 8e) (for further discussion see supplementary material). We note that our primary feather (that the FM is based on), exhibits a considerable built-in pitch-up twist from the base to the tip (see supplementary material Figure S12). Left uncorrected, this would result in excessively high angles of attack and unfavourable flow conditions. Since the FM torque is always directed pitch down, it suggests the torque counteracts the built-in pitch-up twist in the feather (as also suggested in ^4,9^. Furthermore, the change in slope of the torque curve will affect the de-twist rate of the feather, having a higher change with α for higher α and a lower change with α for lower α. Coupled with tuned torsional properties of the feather shaft this behaviour suggests the potential of a passive stabilization mechanism for the α of the feather. To test this hypothesis, we propose a fluid-structure interaction study of the behaviour of the FM, while changing the mechanical properties of the shaft.

The fluid dynamic role of microstructures of biological surfaces have long been recognized, particularly in relation to drag reduction in shark skins ^30^ and dragonfly wings ^23,31^. As in these structures, which also have grooves or valleys, we see a recirculating flow in the spaces between the barbs of the FM (see supplementary material, Figure S9). This type of recirculating flow has been suggested to be the drag-reducing mechanism in ribbed surfaces ^30–32^. However, the riblets in the shark skin are aligned with the flow ^30^ and the valleys on dragonfly wings are aligned transverse to the flow ^31^, while in the feather/FM they are aligned obliquely to the flow, at different directions on the leading and trailing vanes (Figure 2). In a previous study, drag reduction is reduced when riblets are angled relative to the flow rather than aligned or transverse ^30^. Neither the shark skin nor the dragonfly wing has the same shape of the microstructures as the feather/FM. The FM structures are more comparable to blade riblets ^33^, than to the triangular shapes of the dragonfly wing valleys ^29,31^ and concave shape with sharp peaks found in the shark skin structures ^34^. However, the properties of the feather/FM riblets, including the depth to width ratio and thickness to width ratio do not fall within the optimal settings for drag reduction for blade riblets ^30^. Taken together, this suggests relatively low potential for drag reduction of the feather barbs. Despite this, previous studies of feathers have suggested that the ribbed structures of the barbs improve C_d_ ^32,35^, which our findings do not support (i.e. with a higher C_d_ for the FM than for the FEqA, Figure 6). However, the previous studies differ from ours by investigating a secondary flight feather with symmetrical vanes at a location where the feather does not function as an aerofoil on its own, and the flow is directed along the shaft instead of perpendicular to the shaft (e.g. resulting in higher Re). Our results suggest the possibility that the ribbed structure of the feathers may instead result from other factors, such as evolutionary developmental constraints e.g. how the feather grows in the follicle may prevent forming the barbule surface at the dorsal-most position of the barb ^36^. As a result, further studies, varying the height of the barbs above the barbules is required to determine if the ribbed surface indeed improves the aerodynamic performance or if other functions, or lack thereof, provide a better explanation, especially for feathers functioning as aerofoils on their own.

Despite our effort to make the model as accurate a representation of the feather section as possible, we have been forced to make simplifications that may have affected the results and point to potential improvements in future modelling. The most notable difference of the model to the scanned feather relates to how we modelled the barbules (Figure 2). We modelled the barbules as a solid plate with a constant thickness and consequently we note that our model differs in at least four aspects from the real feather: the cross-sectional shape of the space between the barbs, the smoothness of top part of the trailing vane, the porosity of the vanes and the potential difference in the feather in relaxed versus loaded state (see supplementary material for further discussion). These simplifications may affect the flow around the feather and therefore some of the effects seen may not fully represent the behaviour seen in real feathers. However, our FM, as far as we know, represents the hitherto closest model resemblance to a real feather. Real feathers deform under load, but we note that many of the potential effects of flexibility are three dimensional, relying on the deformation of the entire feather, and not just the local section. Therefore, accommodating this deformation requires a larger model than the one we opted for. Still, it would be of interest to incorporate fluid-structure interactions also at this local scale, if nothing else to determine how it differs from the behaviour of the full feather. Nevertheless, a thorough understanding of the basic flow around the feather section, as our model is a first attempt to acquire, is vital for further studies, as well as for understanding the contribution to the aerodynamic forces of each microstructure and their internal relationship.

We opted to only study gliding flight conditions, well aware that birds also flap their wings. Flapping introduces time varying free-stream flow, spanwise pressure gradient and is also likely to increase peak forces (i.e. structural demands). The latter further support the notion that structural properties are a strong driving force in the evolution for optimal morphology of feathers. However, to model flapping reliably requires not only information on how the wing and feather moves, but also how the feather rotates (e.g. pitches) to determine the angle of attack properly. This information is currently not available, to the best of our knowledge, and given the amount of time required for each simulation (~10,000 CPU hours) conducting a parameter search was not feasible and we opted to start by studying gliding flight. Nevertheless, we think our results provide information relevant to natural flight conditions since Jackdaws are capable gliders ^14,37^ and the part of the feather that our model is based on functions as an aerofoil on its own ^1^ and show no fluttering during gliding (LCJ, personal observation). Thus, we consider our results relevant for understanding the aerodynamics and natural selection acting on feathers. None of the model simplifications discussed above are trivial to investigate and will require higher resolution of the model and/or more inclusive modelling, which future studies should yet try to include to investigate their potential impact.

## 5 Conclusion

Our study represents a first step to understand how the detailed morphology of a feather affects aerodynamic forces, with a focus on how our feather section model (FM) performs compared to aerofoils and dragonfly wings at ultra-low Reynolds numbers. Despite a substantial difference in profile shape, the FM exhibited behaviours comparable to aerofoils and cambered plates operating at ultra-low Reynolds numbers. The performance, as displayed in the levels and shape of the C_l_ and C_d_ curves, is affected by the vorticity patterns, camber and flow separation point of the lift generating structure. Our conclusion is that the feather shaft does not negatively affect lift, but since C_d_ of the FM is generally worse compared to the smooth FEqA (Figure 6) the shaft and barbs penalize drag. However, the FM still generates a slightly better C_l_/C_d_ than FEqA. The shaft is the main cause for the drag penalty and has a load carrying function in the feather. This suggests that structural properties of the shaft outweigh the additional drag caused by its obstructive nature on the flow, illustrating an inherent trade-off in the evolutionary development of feather morphology. This and other trade-offs should be considered when seeking inspiration for technical solutions based on feathers.

We find a pitch down torque in the FM, which in combination with the pitch-up pre-twist in the feather is stabilizing. We suggest this helps maintain the force production on the outer part of the feather by passively keeping a favourable α. In addition to resulting in a favourable C_l_/C_d_, keeping the feather in the mid-range α results in a predictable separation point and low amplitude oscillations of the force production see supplementary Figure S8 and Figure S11). The latter likely contributes to better flight control by minimizing overall changes to the wing forces and torques acting on the body during flight. If subsequent testing, incorporating a structural model of the feather, confirms this functionality, it could have significant implications for the micro air vehicle community in their pursuit of designing passive flow control mechanisms.

## 6 Acknowledgments

We are grateful to Kent Persson for discussions regarding the planning of the study, to Stephen Hall for providing the CT-scanning and to Arne Hegemann for providing jackdaws.

## 9 Author contributions

FA contributed to the design, and performed the setup, execution and analysis of the study and wrote the first draft of the manuscript. JR contributed to the design and setup of the study as well as the performance of the solver study. LCJ contributed to the design and analysis of the study. All authors contributed to the final version of the manuscript.

## 10 Data availability statement

The data that support the findings of this study are available from the corresponding author, FA, upon reasonable request.

## 12 Ethical guidelines

Feathers from birds used in this project were obtained through the city of Malmö, Sweden, from pest control, and naturally deceased animals. The CT scanned feather came from the city of Malmö and the additional lower quality scanned feathers came from the latter birds. No animals were euthanized for the purpose of this study, and hence no ethical permission was required.

## References

1. KleinHeerenbrink, M., Johansson, L. C. & Hedenström, A. Multi-cored vortices support function of slotted wing tips of birds in gliding and flapping flight. J. R. Soc. Interface 14, 20170099 (2017).

2. Liu, D. et al. A Brief Review on Aerodynamic Performance of Wingtip Slots and Research Prospect. J. Bionic Eng. 18, 1255–1279 (2021).

3. Feo, T. J., Field, D. J. & Prum, R. O. Barb geometry of asymmetrical feathers reveals a transitional morphology in the evolution of avian flight. Proc. R. Soc. B Biol. Sci. 282, 20142864 (2015).

4. Norberg, R.Å. Function of Vane Asymmetry and Shaft Curvature in Bird Flight Feathers; Inference of Flight Abilit of Archaeopteryx. 16 (1985).

5. Sullivan, T. N., Meyers, M. A. & Arzt, E. Scaling of bird wings and feathers for efficient flight. Sci. Adv. 5, eaat4269 (2019).

6. Eder, H., Fiedler, W. & Pascoe, X. Air-permeable hole-pattern and nose-droop control improve aerodynamic performance of primary feathers. J. Comp. Physiol. A 197, 109–117 (2011).

7. Lucas, A. M. & Stettenheim, P. R. Avian Anatomy Integument Part 1 and 2. (1972).

8. Ennos, A. R., Hickson, J. R. E. & Roberts, A. Functional morphology of the vanes of the fliht feathers of the pegeon cloumba livia. J. Exp. Biol. 198, 1219–1228 (1995).

9. Oorschot, B. K. van, Choroszucha, R. & Tobalske, B. W. Passive aeroelastic deflection of avian primary feathers. Bioinspir. Biomim. 15, 056008 (2020).

10. Terrill, R. S. & Shultz, A. J. Feather function and the evolution of birds. Biol. Rev. 98, 540–566 (2023).

11. Tennekes, H. The Simple Science of Flight, From Insects to Jumbo Jets. (Mit Pr, Cambridge, Mass, 1996).

12. Pennycuick, C. Modelling the Flying Bird. vol. 5 (Elsevier, 2008).

13. Norberg, U. M. Vertebrate Flight. (Springer, 1990).

14. KleinHeerenbrink, M., Warfvinge, K. & Hedenström, A. Wake analysis of aerodynamic components for the glide envelope of a jackdaw (Corvus monedula). J. Exp. Biol. 219, 1572–1581 (2016).

15. Lentink, D. & Kat, R. de. Gliding Swifts Attain Laminar Flow over Rough Wings. PLOS ONE 9, e99901 (2014).

16. Ajanic, E., Paolini, A., Coster, C., Floreano, D. & Johansson, C. Robotic Avian Wing Explains Aerodynamic Advantages of Wing Folding and Stroke Tilting in Flapping Flight. Adv. Intell. Syst. 5, 2200148 (2023).

17. Gasiorek, J. M. K., Jack, L., Douglas, J. F. & Swaffield, J. Fluid Mechanics. (Pearson College Div, Harlow, England; New York, 2006).

18. Klose, B. F., Spedding, G. R. & Jacobs, G. B. Direct numerical simulation of cambered airfoil aerodynamics at Re = 20,000. Preprint at 10.48550/arXiv.2108.04910 (2021).

19. Chen, Z. J., Qin, N. & Nowakowski, A. F. Three-Dimensional Laminar-Separation Bubble on a Cambered Thin Wing at Low Reynolds Numbers. J. Aircr. 50, 152–163 (2013).

20. Winslow, J., Otsuka, H., Govindarajan, B. & Chopra, I. Basic Understanding of Airfoil Characteristics at Low Reynolds Numbers (104–105). J. Aircr. 55, 1050–1061 (2018).

21. Badrya, C., Govindarajan, B. & Chopra, I. Basic Understanding of Unsteady Airfoil Aerodynamics at Low Reynolds Numbers. in 2018 AIAA Aerospace Sciences Meeting (American Institute of Aeronautics and Astronautics, 2018). doi:10.2514/6.2018-2061.

22. Spedding, G. R., Hedenström, A. H., McArthur, J. & Rosén, M. The implications of low-speed fixed-wing aerofoil measurements on the analysis and performance of flapping bird wings. J. Exp. Biol. 211, 215–223 (2008).

23. Okamoto, M., Yasuda, K. & Azuma, A. Aerodynamic characteristics of the wings and body of a dragonfly. J. Exp. Biol. 199, 281–294 (1996).

24. McArthur, J. Aerodynamics of wings at low Reynolds numbers. (University of Southern California, 2007).

25. Tank, J., Smith, L. & Spedding, G. R. On the possibility (or lack thereof) of agreement between experiment and computation of flows over wings at moderate Reynolds number. Interface Focus 7, 20160076 (2017).

26. Hamman, C. W., Klewicki, J. C. & Kirby, R. M. On the Lamb vector divergence in Navier–Stokes flows. J. Fluid Mech. 610, 261–284 (2008).

27. Chang, J., Zhang, Q., He, L. & Zhou, Y. Shedding vortex characteristics analysis of NACA 0012 airfoil at low Reynolds numbers. Energy Rep. 8, 156–174 (2022).

28. Sauvageat, E. Infoscience - EPFL Institutional Repository Viewer. https://infoscience.epfl.ch/entities/publication/d6da3621-0d3e-45a8-ae75-b53ada6d4b02 (2016).

29. Levy, D.-E. & Seifert, A. Simplified dragonfly airfoil aerodynamics at Reynolds numbers below 8000. Phys. Fluids 21, 071901 (2009).

30. Dean, B. & Bhushan, B. Shark-skin surfaces for fluid-drag reduction in turbulent flow: a review. Philos. Trans. R. Soc. Math. Phys. Eng. Sci. 368, 4775–4806 (2010).

31. Kesel, A. B. Aerodynamic characteristics of dragonfly wing sections compared with technical aerofoils. J. Exp. Biol. 203, 3125–3135 (2000).

32. Chen, H., Rao, F., Shang, X., Zhang, D. & Hagiwara, I. Biomimetic Drag Reduction Study on Herringbone Riblets of Bird Feather. J. Bionic Eng. 10, 341–349 (2013).

33. Bechert, D. W., Bruse, M., Hage, W., Hoeven, J.G.T.V.D. & Hoppe, G. Experiments on drag-reducing surfaces and their optimization with an adjustable geometry. J. Fluid Mech. 338, 59–87 (1997).

34. Oeffner, J. & Lauder, G. V. The hydrodynamic function of shark skin and two biomimetic applications. J. Exp. Biol. 215, 785–795 (2012).

35. Benschop, H. O. G. & Breugem, W.-P. Drag reduction by herringbone riblet texture in direct numerical simulations of turbulent channel flow. J. Turbul. 18, 717–759 (2017).

36. Prum, R. O. Development and evolutionary origin of feathers. J. Exp. Zool. 285, 291–306 (1999).

37. Rosén, M. & Hedenström, A. Gliding flight in a jackdaw: a wind tunnel study. J. Exp. Biol. 204, 1153–1166 (2001).

