## Supplemental figures and text for "Feather aerodynamics suggest importance of lift and flow predictability over drag minimization"

**Summary of content**
This document contains supplementary figures and text. The material is divided into four sections, with the first section containing figures referenced from the main document, the second section containing a video, the third section added details of the results and discussion, and finally additional references used in the supplement.

### *I. Figure list*

Figure S1. Measurements of the feather
Figure S2 CAD model vs CT scan, profile matchup for front vane
Figure S3 Domain and mesh
Figure S4. Mesh Sensitivity, mesh 1, mesh 2 (FM) and mesh 3
Figure S5. Results of the numerical validation study
Figure S6. Results of extended aerofoil investigation
Figure S7. Performance of the FM varies over time
Figure S8. Separation point, along the chord, of flow on top side for FM and FEqA
Figure S9 Vector flow field on the leading vane and shaft area.
Figure S10 Cl curves and time averages for FEqA
Figure S11 Shedding amplitudes for FM, dragonfly like foil and Eppler-E61
Figure S12 Shape and twist of the feather

### *II. Movie List*

Movie M1 In this document, still image of shedding video. See separate file for video.

### *III. Additional details*

**Results and discussion**
Separation bubble
Torque pattern
Feather representation

### *IV. References*

### I. Figures

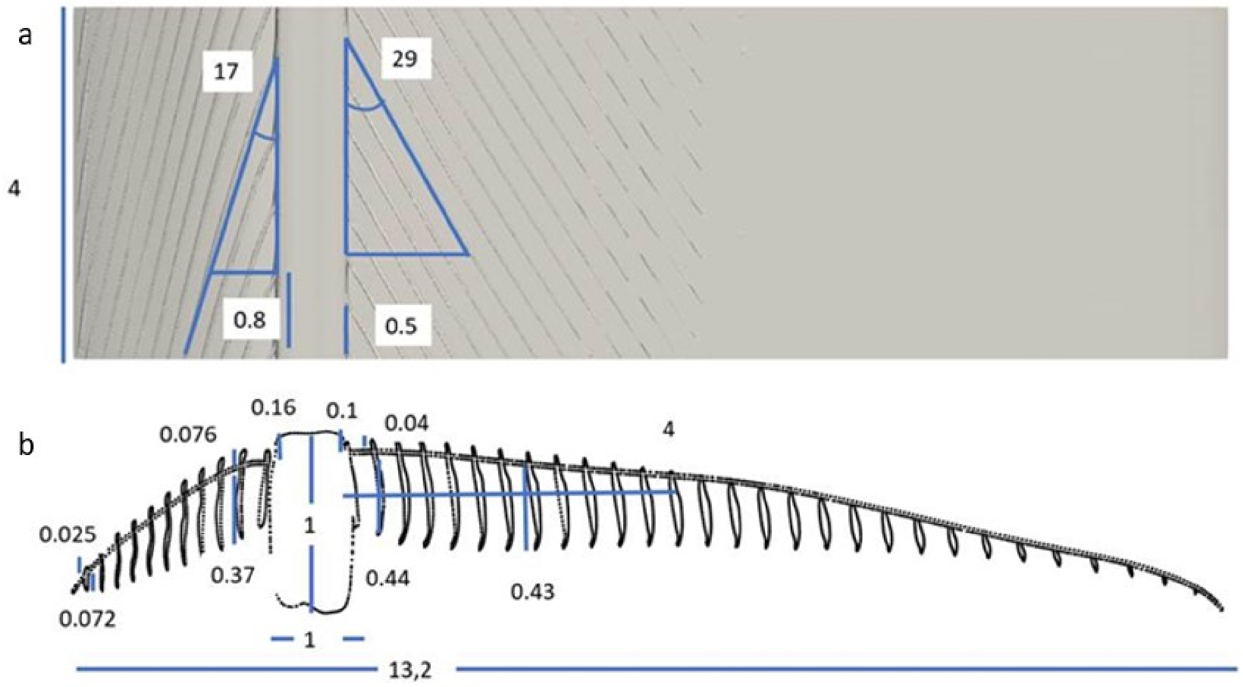


**Figure S1.** Measurements of the feather. *(a)* Width of the model as well as angle of barbs of front and back vane and the distance between the barbs. *(b)* Height and width of shaft and barbs, distance from shaft to end of barbs on top surface on back vane as well as the chord length. Note that the scales in length and height are not the same. All measurements in mm and degrees.

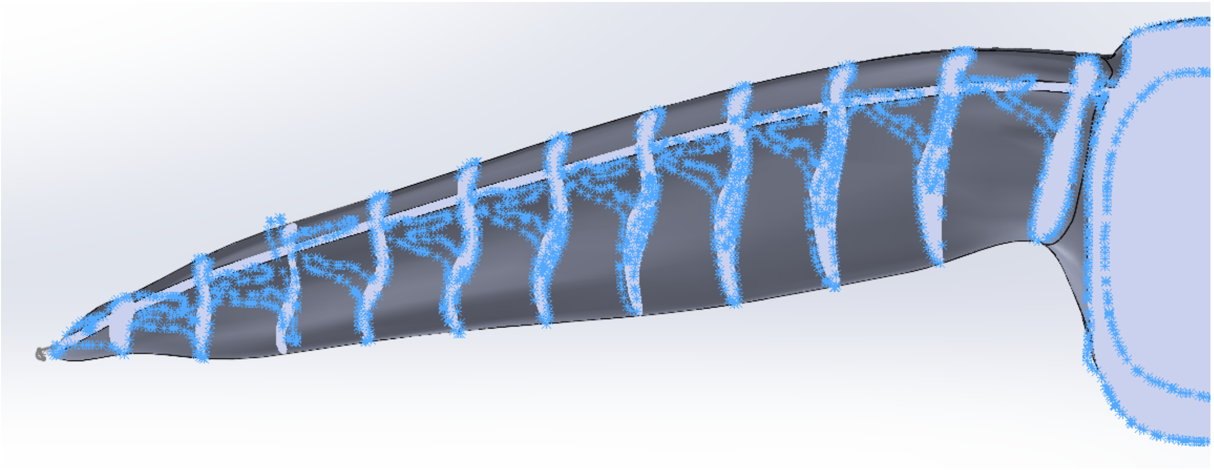

**Figure S2.** CAD model in relation to CT-scan. The grey background object shows the CAD-model of the FM, and the blue asterisks represent the cross section of the feather. Cross sections of the CAD barbs (light grey) and shaft match the scan well, but rotated and displaced barbs were corrected in the model. The barbule plane is matched to the height of the proximal barbules.

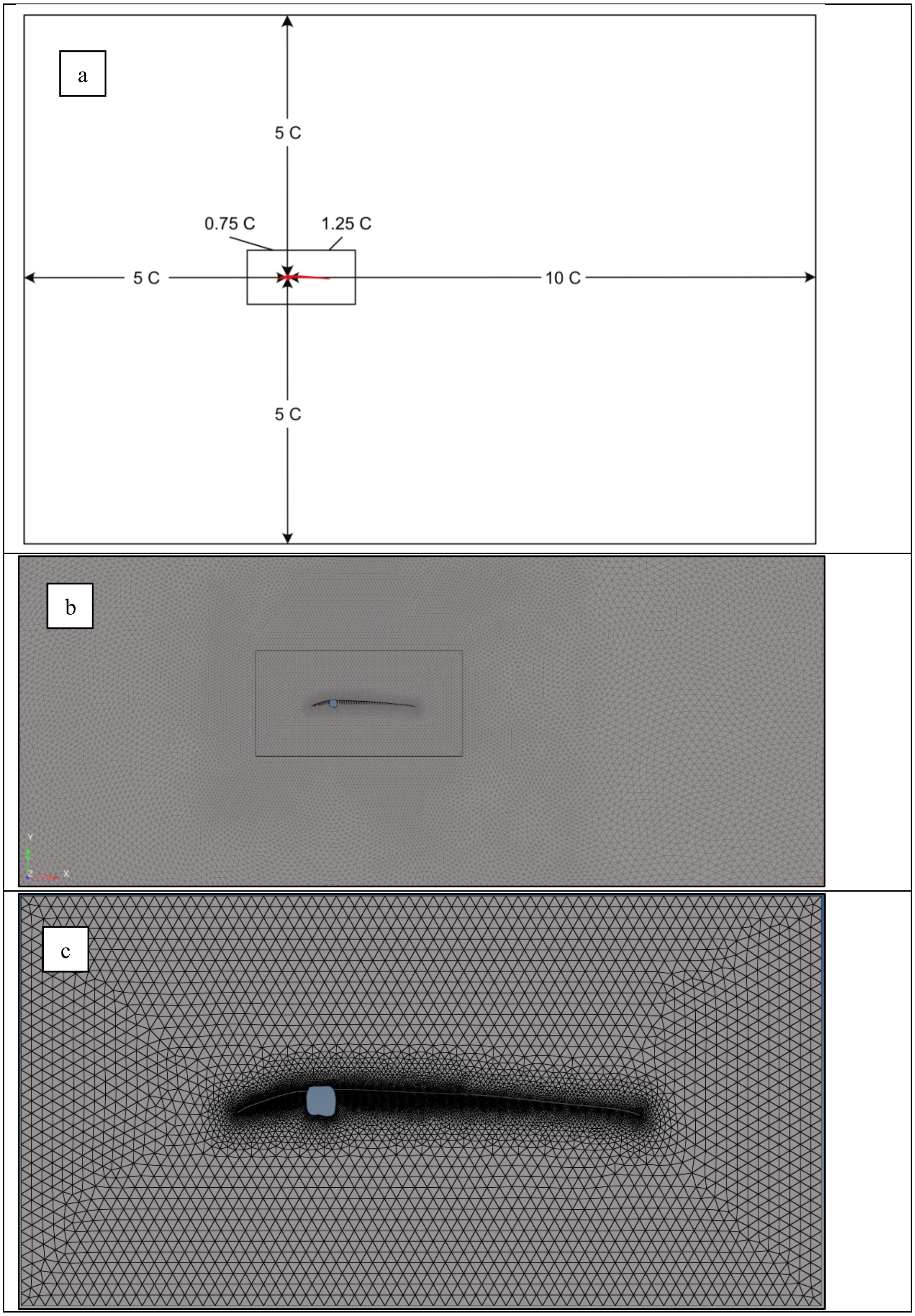


Figure S3. Mesh and domain size. *(a)* Domain size, FM in the middle surrounded by the inner box, sized 2x1 c (where c is the chord of the model), and the outer domain size 15x10c. *(b)* Part of the mesh domain with Inner part of mesh domain enlarged in *(c)*.

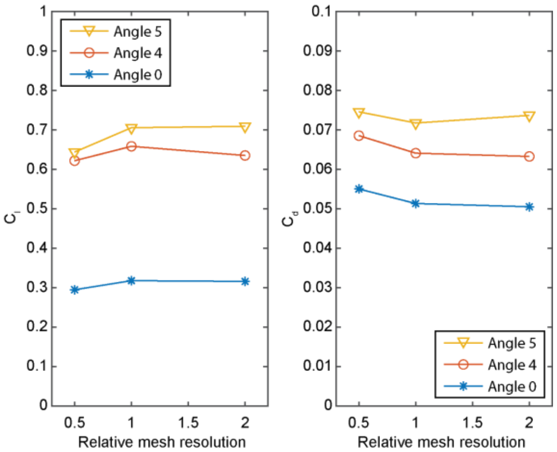


**Figure S4**. Mesh sensitivity analysis, where mesh 1 has half the resolution of mesh 2 (FM) and mesh 3 has twice the resolution of mesh 2. Mesh 1 shows the most difference from the other two and the $C_l$ deviation between mesh 2 (FM) to mesh 3 was less than 1 percent for α=0.41° and α=5.41° and 3% for α=4.41°, the $C_d$ deviation was less 2% for α=0.41° and α=4.41 and less 3% for α=5.41°.

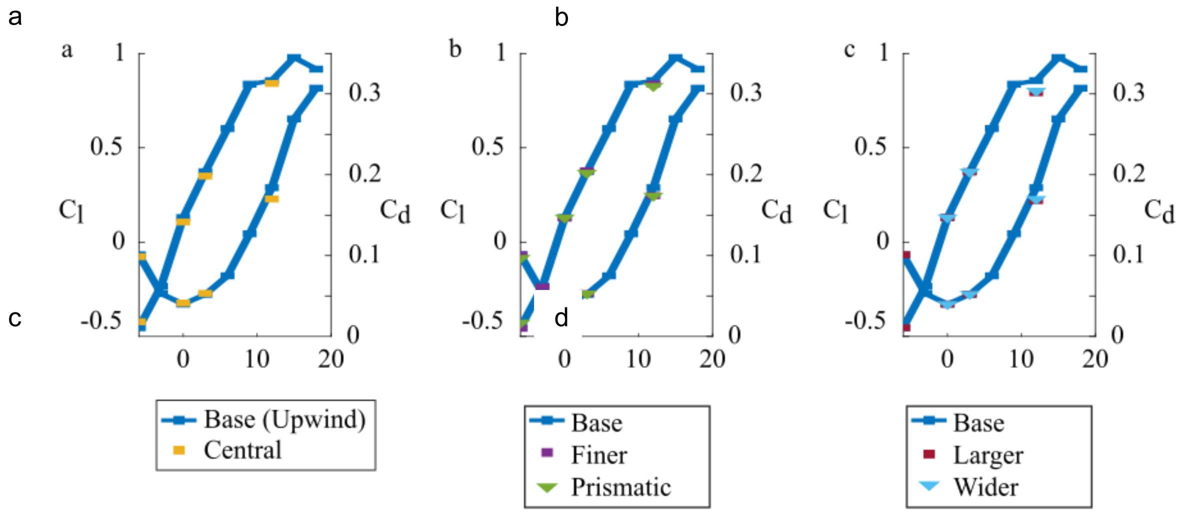


**Figure S5**. Results of a numerical validation study using an Eppler (E387) foil in OpenFoam. Any modification of the modelling settings (convective scheme *(a)*, mesh resolution *(b)*, and domain size *(c)*) show no deviations from the base setting showing a robustness in the simulation approach. All models were run at Re = 10 000.

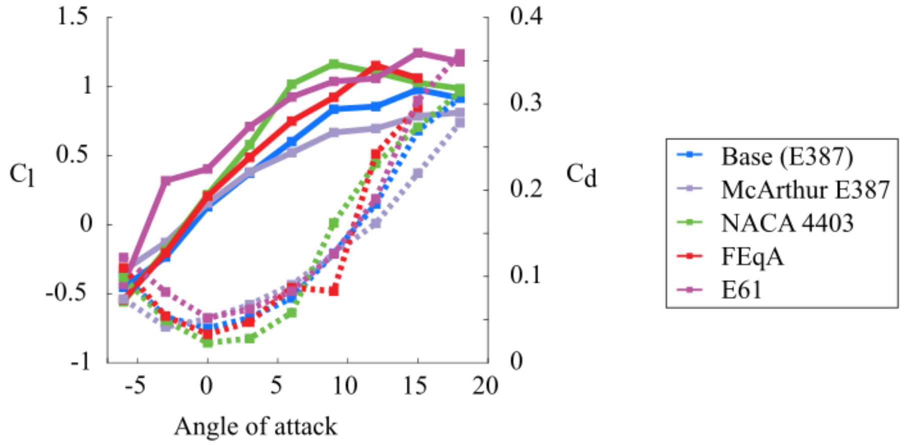


**Figure S6**. Results of extended aerofoil investigation modelling an Eppler E387 and E61, the NACA4403 and FEqA, shown together with experimental data from E387 [1] .The relationship between the simulated aerofoils is similar to the results presented in Figure 6, supporting the notion that the E387 experimental data stands out.

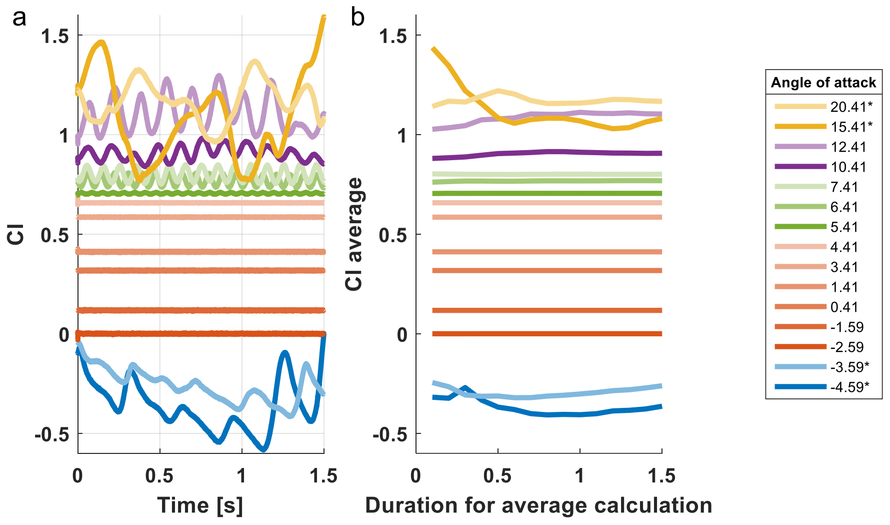


**Figure S7.** Performance of the FM varies over time. *(a)* The lift coefficient (Cl) during the last 1.5 seconds of the simulation, where color groups represent different fluctuation patterns in the lift coefficient, group 1=blue, group 2= orange, group 3=green, group 4=purple and group 5=yellow. Non-periodic fluctuations in groups 1 and 5 may be due to insufficient running time for the model or be due to chaotic lift generation [2]. *(b)* The time average lift coefficients, with x-axis showing the time duration used for the calculation. For example, x =1 represents an average over the time interval 0.5-1.5 in panel a. Asterixis in legend mark angle of attacks where the mean data differs more than 2.5% between an average of 1.5 and 1.4 second. These latter data are kept to indicate trends in the results, but should be interpreted with caution.

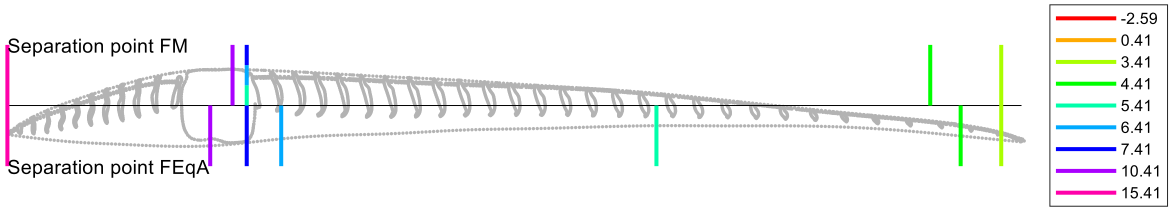


**Figure S8.** Separation point along the chord of the flow <u>on the top side</u> for FM (above black line) and FEqA (below black line). Note that the shaft has a strong influence on the point of separation for the FM.

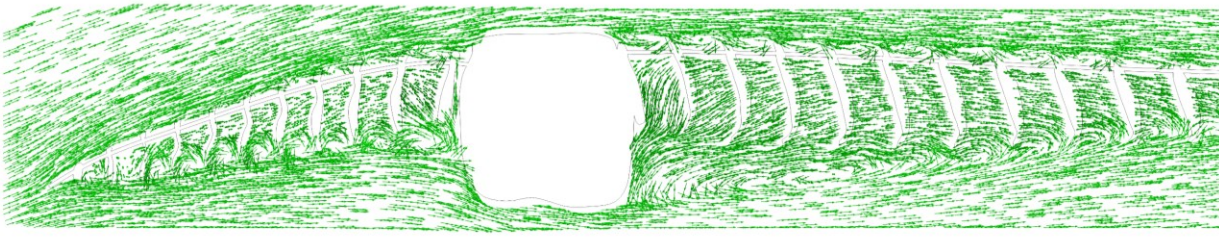

**Figure S9.** Flow vectors illustrating rotational structures for the front half of the FM in a single plane. On the front vane we see rotation between the barbs at the bottom side of the vane, but also on the top side. The stagnation point on the shaft is located at ~1/4 from the bottom of the shaft. On the trailing vane we see a large rotating structure behind the shaft and small rotational structures between the barbs on the top surface. Angle of attack is α=4.41.

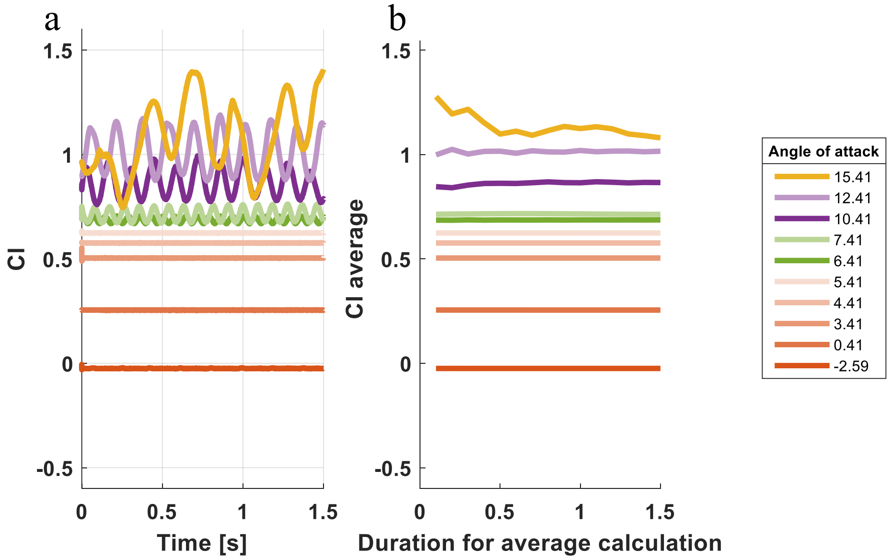


**Figure S10.** Performance of the FEqA varies over time. *(a)* The lift coefficient (Cl) during the last 1.5 seconds of the simulation, where color groups represent different fluctuation patterns in the lift coefficient, group 2= orange, group 3=green, group 4=purple and group 5=yellow. *(b)* The time average lift coefficients, with x-axis showing the time used for the calculation. For example, x =1 represents an average of time interval 0.5-1.5 in panel a.

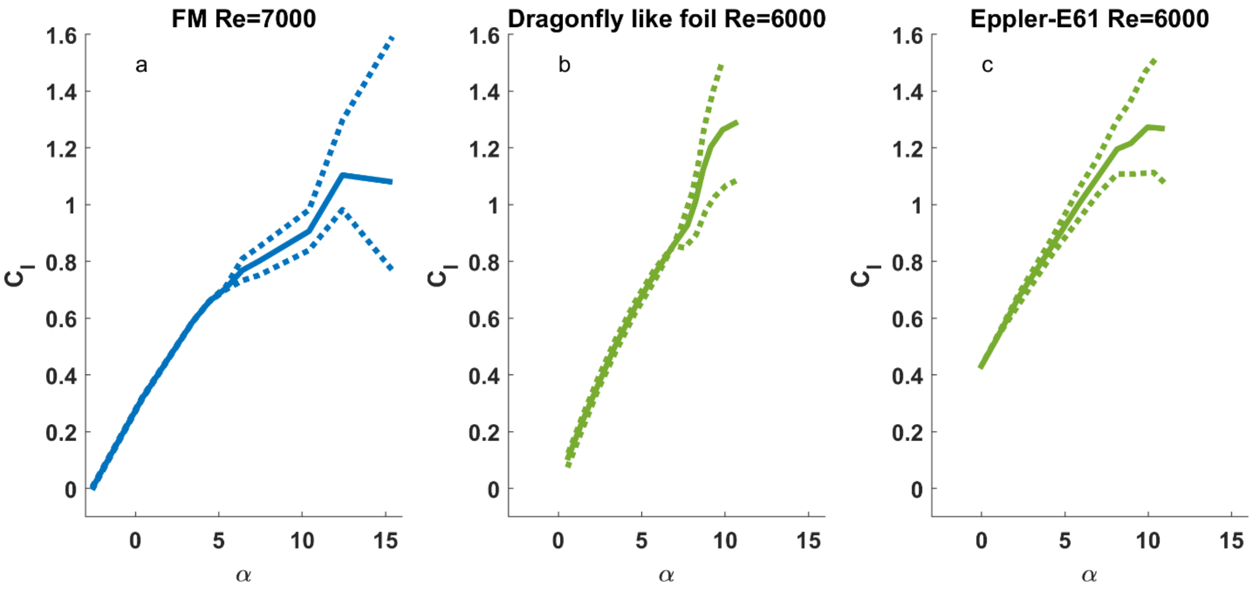


**Figure S11.** The mean lift coefficient (solid line), with dotted lines indicating the variation (max/min values) across angles of attack (α), for *(a)* FM, *(b)* dragon fly like wing and *(c)* Eppler-E61 aerofoils. For the Eppler foil (c) variation increases continuously as angle of attack increases, while for FM (a) and the dragonfly wing (b) variation remains low and then increases at a critical α of ~6-8 degrees. This suggests more stable lift production at low α for the biologically inspired foils than for the Eppler foil. Data for the FM is the mean during the last 1.5 s of simulation, while b) and c) are adapted from [3].

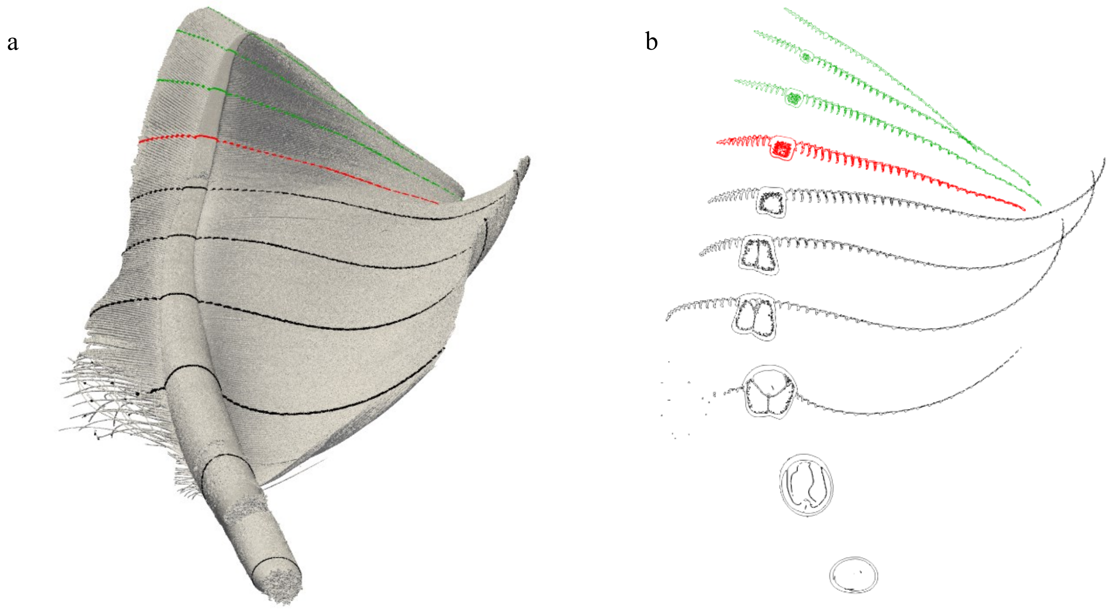


**Figure S12.** The shape and twist of the feather changes along its length. (a) CT scanned feather with several cross sections marked in black (proximal part with overlap from adjacent feather), red (cross section used for model) and green (distal part). (b) Same cross sections as in (a) showing the shape of the shaft changes from round to square back to round again when going from base to tip. The middle of the shaft is changing from hollow to different chambers filled with foam (medulla) to solid at the tip. The tip of the feather, which acts as an aerofoil on its own, has a pitch up twist

**Videos**

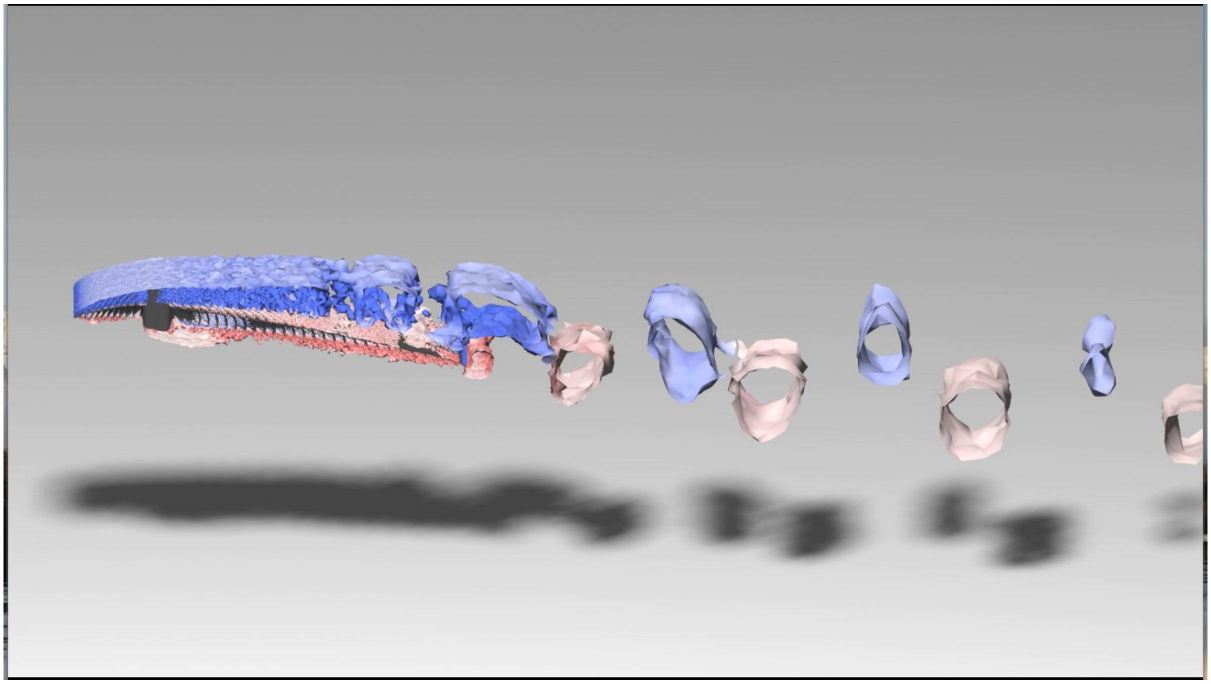

**Movie M1.** Shedding patterns for α=7.41, illustrated as surfaces of Q-criterion at level 50, for one oscillation cycle. See separate file.

### II. Additional details

#### Results and discussion

##### Separation bubble

At low Re and relevant α one separation structure often seen on airfoils is the laminar separation bubble [4,5]. However, we did not detect a laminar separation bubble in our simulations. There are several geometrical differences between the airfoils used and the FM, where the FM, for example, has a thinner leading edge, a lower curvature on the upper surface as well as the thickest part of the profile relatively closer to the leading edge. In addition, the FM has barbs which breaks the smooth surface otherwise common in airfoils, although these barbs do not seem to impact the flow's ability to stay aligned with the surface until after the shaft or at high α where a laminar separation bubble is not expected. All of the mentioned geometrical features have the potential to affect the flow close to the FM and thereby affect the formation of a laminar separation bubble. Further studies are required to determine which of the factors is most important and if the lack of a laminar separation bubble can e.g. explain the low variation in the lift production at low angles of attack (Fig. S10).

##### Torque pattern

The movement of center of pressure has one outlier for $5.41 \leq \alpha \leq 10.41$, not fitting the pattern (Fig. 8f). When comparing the FM and the FEqA which does not have the same pronounced outlier a difference can be found. The flow separation point (Fig. S3) is moved from the trailing edge to the shaft and is then fixed at the trailing end of the shaft for midrange α for the FM, whilst the separation point is continuously moving forwards for each α on the FEaA. Additionally, the lower lift coefficient for the FEqA could be the reason for its lower moment compared to the FM. However, it is also possible that this results from a combination of lower lift and a shorter moment arm since the center of pressure is closer to the shaft than for the FM.

##### Feather representation

There are four main aspects where our FM model differs from the real feather: the cross-sectional shape of the space between the barbs, the smoothness of top part of the trailing vane, the porosity of the vanes and the potential difference in the feather in relaxed versus loaded state. First, the cross-sectional shape of the space between the barbes become square instead of triangular on the bottom side (see supplementary material, Fig. S2). This may affect the size and stability of the vortices formed in the openings of the space between the barbs. However, this primarily affects the space above the vortices, where the velocity is low, reducing the potential impact. Second, the trailing half of the trailing vane is smooth on the top surface since the barbs do not protrude above the surface (Fig. 2). This may affect the behavior of the boundary layer at the second half of the trailing vane. One potential effect could be an earlier separation of the flow at the top surface of the FM, although we, given the size of the structures, find it unlikely they would have a major impact on the results. Regardless, future studies should try to explore the effect of including barbs protruding the barbule plane also at the trailing half of the vane. Third, the barbule plane is not porous, as has been demonstrated for real feathers [6,7], but due to the range of the recorded data and the methods used, we were not convinced it would be appropriate to include it in our model. Feather porosity has been measured through the barbule plane and also been suggested to result from pores forming close to the shaft [6,7]. Porosity in vanes is partly determined by the spacing of the barbs [8] and the barbules [7]. The porosity of feathers also depends on where it is measured, which species of bird is used and which feather and vane is tested [6,7]. Given the large range of the permeability found in the literature and no direct link between permeability and morphological measurements, we did not consider it possible to estimate the permeability of the jackdaw feather. In addition to permeability of the vanes, pores close to the shaft have been suggested on the basis of microscopic studies of feathers [6]. These pores have been suggested to improve the aerodynamic performance of the feather since closing the pores with water-soluble gel resulted in worse performance [6]. We did not include pores in this study since we did not find support for them in the CT-scan. Also, given that the use of gel to close the pores could have resulted in changes in the mechanical properties of the feathers, which could have affected the findings we were not convinced it would be appropriate to include them. Fourth, there is an overarching uncertainty of our model due to the potential difference of the feather shape in a relaxed and loaded state and the resolution capacity of the fluid dynamic simulations. We have modelled the feather section as a solid structure, while the flexibility of real feathers is likely to result in deformations subject to the aerodynamic forces, especially at the trailing vane, which is less stiff than the leading vane. This suggests that some of the behavior that we see in the FM aerodynamics may not fully represent the behavior seen in real feathers. We find it likely that the elastic deformations of the feather may modify camber, but also the shape of the trailing edge of the feather/FM and hence the shedding behavior and potentially the peak $C_l/C_d$. However, given that we do not know how a primary feather would deform we opted for this simplification in this initial study, but would suggest that future studies may benefit from including structural modelling.

### III.References


1. McArthur, J. Aerodynamics of wings at low Reynolds numbers. (University of Southern California, 2007).
2. Chang, J., Zhang, Q., He, L. & Zhou, Y. Shedding vortex characteristics analysis of NACA 0012 airfoil at low Reynolds numbers. *Energy Reports* **8**, 156–174 (2022).
3. Levy, D.-E. & Seifert, A. Simplified dragonfly airfoil aerodynamics at Reynolds numbers below 8000. *Physics of Fluids* **21**, 071901 (2009).
4. Klose, B. F., Spedding, G. R. & Jacobs, G. B. Direct numerical simulation of cambered airfoil aerodynamics at Re = 20,000. Preprint at https://doi.org/10.48550/arXiv.2108.04910 (2021).
5. Winslow, J., Otsuka, H., Govindarajan, B. & Chopra, I. Basic Understanding of Airfoil Characteristics at Low Reynolds Numbers (104–105). *Journal of Aircraft* **55**, 1050–1061 (2018).
6. Eder, H., Fiedler, W. & Pascoe, X. Air-permeable hole-pattern and nose-droop control improve aerodynamic performance of primary feathers. *J Comp Physiol A* **197**, 109–117 (2011).
7. Müller, W. & Patone, G. Air transmissivity of feathers. *Journal of Experimental Biology* **201**, 2591–2599 (1998).
8. Lucas, A. M. & Stettenheim, P. R. *Avian Anatomy Integument Part 1 and 2*. (1972).
